# Asymmetric expression of homoeologous genes in wheat roots modulates the early phase of iron-deficiency signalling

**DOI:** 10.1101/2022.08.19.504540

**Authors:** Gazaldeep Kaur, Varsha Meena, Anil Kumar, Gaurav Suman, Deepshikha Tyagi, Riya Joon, Janneke Balk, Ajay K Pandey

**Affiliations:** National Agri-Food Biotechnology Institute (Department of Biotechnology), Sector 81, Knowledge City, S.A.S. Nagar, Mohali-140306, Punjab, India; Regional Centre for Biotechnology, Faridabad 121001, India; Department of Biochemistry and Metabolism, John Innes Centre, Colney Lane, Norwich, NR4 7 UH, United Kingdom; School of Biological Sciences, University of East Anglia, Norwich, NR4 7TJ, United Kingdom

**Keywords:** iron, hexaploid wheat, early response, iron deficiency, iron transport, transcription factor, yellow stripe-like transporter, genome-bias, polyploidy

## Abstract

Iron (Fe) limitation leads to dramatic changes in gene expression in plants, to induce iron uptake and mobilization, while at the same time restrict Fe-dependent metabolism and growth. Although transcriptional changes in response to Fe deficiency have recently been charted in wheat, this was performed at a stage when photosynthesis and growth were severely impacted, confounding primary and secondary responses. Here, we specifically uncover the transcriptional landscape of wheat roots during the early stages of the Fe deficiency response (4 and 8 days) and after Fe resupply. Root growth was significantly inhibited at day 4, but chlorosis only became apparent on day 8. The number of differentially expressed genes increased from 1386 on day 4 to 3538 on day 8, with an overlap of 2006 genes. Genes with dynamic changes in expression patterns include membrane transporters and transcription factors shown to be involved in Fe homeostasis in other plant species. Comparative analysis of the Fe deficiency response at 4, 8 and 20 days identified a core set of Fe-regulated genes.

Analysis of the expression of homoeologs suggests an increase in induction bias at 8 days compared to 4 days particularly, A genome contributing high at 4 days and the A+D genomes at 8 days. Overall, our work will contribute towards fundamental knowledge of the Fe signalling networks in wheat and point to the interplay of the three sub-genomes in this hexaploid species to fine tune the transcriptional response.

## 1. Introduction

Iron (Fe) is an essential micronutrient involved in primary metabolic processes such as photosynthesis, respiration, nitrogen and sulphur assimilation, chlorophyll biosynthesis, DNA synthesis and many more catabolic and anabolic pathways (Briat et al., 2015; Przybyla- Toscano et al., 2021). Fe is an abundant element in the geosphere, but largely unavailable in its oxidized, ferric form, Fe(III). As a consequence, higher plants have developed high-affinity Fe uptake mechanisms that are engaged when the demand for iron for photosynthesis and growth is larger than the supply through the roots (Connorton & Balk, 2019; Zhang et al., 2019).

Dicots or non-graminaceous plants generally follow the strategy I mode of Fe uptake, also referred to as reduction-based strategy, involving reduction of Fe(III) to ferrous iron, Fe(II), by the ferric chelate reductase FRO2. Fe(II) is then taken up by the iron regulated transporter IRT1. FRO2 and IRT1 work in concert with a proton extrusion pump (H+ATPase, AHA2) which increases the solubility of Fe (III) complexes (Martín-Barranco et al., 2020). Monocots including graminaceous plants secrete specific metabolites into the rhizosphere to facilitate Fe uptake, known as the strategy II mode. The Fe-chelating metabolites are generically called phytosiderophores (PS). The PS are exported by the transporter of mugineic acid (TOM) and the Fe-PS complexes are imported into root cells by the high-affinity yellow stripe-like transporter YSL (Curie et al., 2001; Inoue et al., 2009; Yordem et al., 2011). The YSL proteins are a subfamily of the oligopeptide transporter family (OPT), encoding integral membrane proteins with multi-substrate transport functions. The YSL transporters in plants are involved in iron transport from either the extracellular environment or from chloroplasts (Kumar et al., 2019; Lubkowitz, 2011). The regulation of Fe uptake is primarily at the transcriptional level, and is well studied in Arabidopsis and rice (Gao et al., 2019). Multiple studies have elucidated the crucial role of basic helix-loop-helix (bHLH) and MYB transcription factors (TF) in the maintenance of Fe homeostasis (Wang et al., 2019; Wu & Ling, 2019). Interestingly, the bHLH transcription cascade, including iron-binding E3 ligases that regulate key bHLH TFs post-translationally, are conserved between dicots and monocots.

Genes involved in Fe homeostasis have been utilized in biotechnology approaches to increase the Fe concentration in crops, especially in rice and cassava(Masuda et al., 2013; Narayanan et al., 2019). In wheat, the third most cultivated crop and a staple food for major parts of the world, overexpression of the wheat vacuolar iron transporter VIT2 in the endosperm, or constitutive overexpression of nicotianamine synthase (NAS2) from rice have been shown to improve the iron and zinc concentration of wheat flours (Beasley et al., 2019; Connorton et al., 2017; Harrington et al., 2022; Singh et al., 2017). To identify other genes that can be targeted for biofortification, transcriptome analysis of the Fe deficiency response in wheat can be an important tool to (i) identify paralogues of multi-gene families that are involved in Fe homeostasis and, (ii) establish whether one particular homoeolog of a gene triad in hexaploid bread wheat plays a dominant role. This knowledge will be crucial for gene- editing strategies to change the function of endogenous genes.

Time-course experiments have aimed to separate early from late and/or indirect responses to Fe deficiency in model organisms such as the green alga Chlamydomonas and Arabidopsis (Urzica et al., 2012) and in crops such as soybean and barley(Atencio et al., 2021; Darbani et al., 2013). Previous studies in wheat to identify Fe-regulated genes focussed on the late stages of Fe deficiency when the plants displayed severe adjustments to Fe limitation and secondary physiological responses(Kaur et al., 2019; Wang et al., 2019). However, the molecular events leading to the early adaption to the Fe deficiency remain unaddressed.

In the present work, our goal was to decipher the early transcriptional responses to Fe deficiency in wheat roots, in particular responses that are reverted by resupply of Fe. We performed RNAseq analysis at 4 and 8 days of Fe deficiency and 2 days after resupply of the Fe. Our data reveal new insights in differentially expressed genes and help categorize multiple iron homeostasis genes specific to the early pre-symptomatic (4 days) and symptomatic stages (8 days) of Fe deficiency, including regulatory genes and genome-biased expression of homoeolog genes.

## 2. Materials and methods

### 2.1 Plant materials, experimental design and leaf chlorophyll measurement

Hexaploid bread wheat, variety ‘C-306’, was grown as described earlier (Kaur et al., 2019) with minor modifications. Seeds were surface sterilized using 1.2% (w/v) sodium hypochlorite in 10% (v/v) ethanol for 3 min, rinsed twice with sterile milliQ water, and then kept on wet filter paper at 4 °C overnight. Sterilized seeds were germinated at room temperature for 5 days. Uniform healthy seedlings were transferred to a hydroponic system with Hoagland media containing: 6 mM KNO_3_, 1 mM MgSO_4_.7H_2_O, 2 mM Ca(NO_3_).4H_2_O, 0.2 mM KH_2_PO_4_, 80 µM Fe (III) EDTA, 0.25 mM H_3_BO_3_, 2 µM MnSO_4_.H_2_O, 2µM ZnSO_4_.7H_2_0, 0.5 µM CuSO_4_.5H_2_0, 0.5 µM Na_2_.MoO_4_ and 50 µM KCl. After 5 days of synchronized growth seedlings were exposed to Fe deficiency (2 µM of Fe (III) EDTA) or continued to growth under Fe sufficiency (80 µM Fe (III) EDTA). Tissue samples were collected after 4 and 8 days of treatment. For the Fe resupply treatment, the Fe deficient medium was replaced by medium containing 80 µM Fe (III) EDTA for an additional 2 days to allow recovery from Fe stress. All these experiments were carried out in a growth chamber under controlled environmental conditions at 22 °C temperature, at a photoperiod of 16 h day and 8 h night, 65%–70% humidity and 300 nm photon m^−2^s^−2^of light. The experiments were performed in triplicate, with each replicate consisting of twelve seedlings. At the mentioned time points, the seedlings or the tissue samples were rapidly frozen in liquid nitrogen.

### 2.2 Measurement of chlorophyll content

The total chlorophyll content was expressed in terms of SPAD index. The SPAD index was determined on the fully expanded youngest leaves (middle of the long axis) in 3-4 positions on nine individual plants using a portable chlorophyll meter (Konica Minolta SPAD-502) and mean value was calculated. These experiments were repeated three times with similar observations.

### 2.3 RNA extraction and transcriptome analysis

Total RNA from root tissues was extracted using TRIzol® Reagent (Invitrogen™, USA). Genomic DNA was removed using the TURBO™ DNase kit (Ambion, Life Technologies). The integrity of the RNA samples (RIN≥8.5) was checked using a Bioanalyzer (Agilent, USA) prior to library preparation. Paired-ended libraries were prepared using the Illumina TruSeq stranded mRNA library prep kit as per the instructions (Illumina Inc., USA) by Eurofins, Bangalore, India.

Analysis of the raw sequencing data was done in a stepwise manner as described earlier (Kaur et al., 2019) and Supplementary Figure S1. The sequence reads were trimmed using Trimmomatic -0.35 to remove adapters and processed further to obtain high-quality clean reads with a minimum length filter of 100bp. The high-quality (QV>20), paired-end reads were used for reference-based read mapping. The transcriptome of *Triticum aestivum* L. from EnsemblPlants (IWGSC RefSeq v1.0, ensembl release 47) was taken as a reference for the analysis. The obtained reads were pseudo-mapped against the reference sequences to get raw counts using Kallisto followed by gene-level abundance summarization using the tximport package in R. Normalized expression counts were obtained through DESeq2. Principal Component Analysis (PCA) was performed using the VST transformed expression counts for the 500 genes with the highest variation across conditions and plotted using ggplot2 to observe the closeness among replicates and deficient/resupply conditions. Differential expression analysis for the conditions *w*.*r*.*t*. control samples was performed using DESeq2 wherein genes with Log2 Fold Change (LFC)>1 or <1 with a corrected p-value <0.05 were considered to be significantly differentially expressed (DE). For the construction of clustered dendrogram of DEGs according to their expression patterns, hierarchical clustering was performed using hclust package in R, nbclust package was used to determine the number of clusters for the dendrogram.

### 2.4 Functional annotation and enrichment analysis of DEGs

Comprehensive gene annotation of wheat sequences was done using KOBAS 3.0 (Xie et al., 2011). Further, annotate module of *Oryza sativa* sequences as (BLASTP, cutoff 1e-5), in addition to the wheat RefSeq v1.0 annotation was used for predicting gene function. MapMan was used to visualize the pathways involving wheat differentially expressed genes (DEGs). Pathway enrichment analysis was performed using KOBAS standalone tool. Functional enrichment analysis was performed using Panther (reference: PMC6519453), through the gene ontology resource (http://geneontology.org/, latest release: 2021-08-18). Significantly enriched GO terms were identified through Fisher’s exact test and Bonferroni correction for multiple testing. MeV and gplots package were used to construct heatmaps for the selected DEGs using the normalized expression values of genes. The data generated from this study has been deposited in the NCBI Sequence Read Archive (SRA) database (Submission ID- SUB10305760 and BioProjectID- PRJNA759756).

### 2.5 Analysis of wheat homoeolog expression and induction bias

Specific changes were made to the analysis pipeline to increase the sensitivity so as to differentiate among different homoeologs and correctly quantify them. As mentioned earlier the tophat2-cufflinks pipeline was used to obtain normalized expression counts for the “accepted triads” (Kaur et al., 2019).These triads are single and closely related copies of a gene on each of the three sub-genomes. Based on these specifications, 12332, 12312, 12488, and 12295 triads were considered for expression bias and induction studies for 4 and 8 days of Fe deficiency and the respective resupply time points. Triads showing a biased induction (up-or down-regulation of 2-fold, with a corrected p-value<0.05) from a sub-genome under Fe were categorized accordingly.

## 3. Results

### 3.1 Physiological changes at the early phase of Fe deficiency

To examine early changes in gene expression in response to Fe deficiency, 5-day-old wheat seedlings were transferred in hydroponic medium to expose them to Fe deficiency (2 µM Fe EDTA) or maintain them at Fe sufficiency (80 µM Fe EDTA). Roots and shoots were harvested after 4 days and 8 days (Figure 1A). At the phenotypic level, a significant decrease in root length was observed following 4 days of Fe deficiency and this phenotype was even more pronounced on day 8 (Figure 1B and C). After 4 days of Fe deficiency, no significant changes in chlorophyll were observed as reflected in similar SPAD index in control and Fe deficient seedlings. A significant decrease in the SPAD index (16%) was observed at day 8 of Fe deficiency, with visible chlorosis compared to control seedlings (Figure 1D). Although no effect on shoot or shoot biomass was observed at 4 days, a significant reduction was observed at 8 days of Fe deficiency (Figure 1E). The root and shoot biomass was significantly decreased at 8 days post Fe deficiency (Figure 1E). To distinguish general changes in gene expression (e.g. because of growth adaptation) from Fe-specific changes, the Fe-deficient wheat seedlings were subsequently replenished with Fe for two additional days to rescue the partial phenotype. Thus, analysis of plant growth and chlorophyll confirmed that the 4- and 8-day time points of Fe deficiency represent the early stages of the phenotypic response, and were used, along with the resupply treatments, to study comparative gene expression changes in the roots.

**Figure 1:**
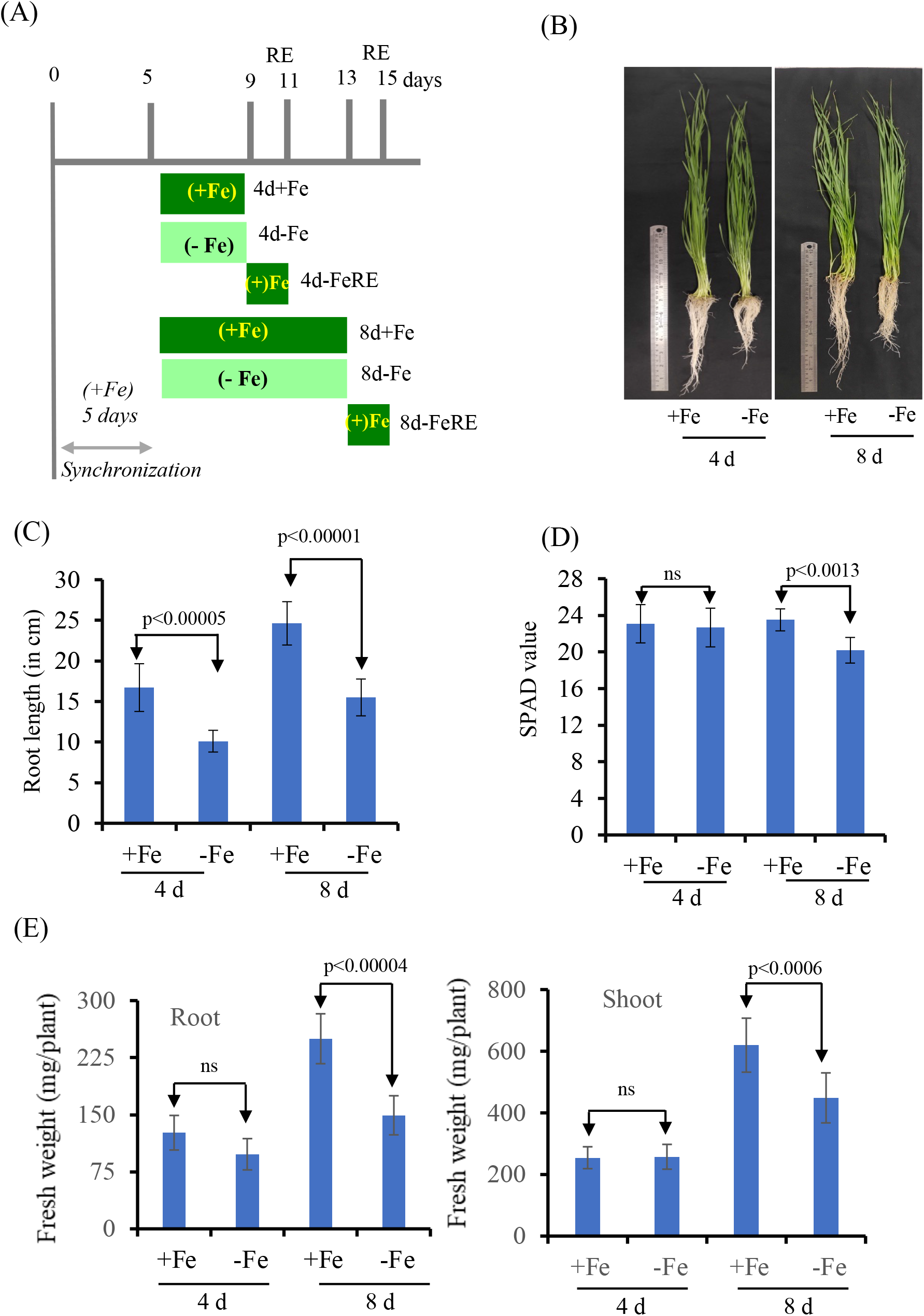
Phenotypic effect of short-term Fe deficiency in hydroponic growth conditions. **(A)** Five-day-old wheat seedlings grown with sufficient Fe (80 µM, labelled +Fe) were subjected to Fe deficiency (2 µM Fe, labelled -Fe) for 4 d and 8 d. Control seedlings were grown for the same length of time under Fe sufficiency. For later RNAseq analysis, seedlings grown -Fe were transferred to Fe-sufficient medium and grown for another 2 days (re-supply, labelled RE). (B) Phenotypic changes in the wheat seedlings under Fe-deficiency condition. (C) Root length (D) Chlorophyll content by SPAD analysis. (E) Fresh weight of roots and shoots (mg/plant). Values at the indicated days of treatment are given as averages ± SD (n = 12).

#### Changes in the number of DEGs in roots during Fe deficiency and resupply

Comparative analysis of RNAseq data revealed distinct gene expression profiles at each of the Fe treatments and time points. PCA analysis confirmed good repeatability clustering of the biological replicates with each other (Supplementary Figure S2). Based on the first principal component, transcriptomes for the 4- and 8-day time points were found to be clustered together for the respective controls and deficiency conditions. Relatively higher variance was observed after Fe resupply between the 4- and 8-days treatments. The transcriptome of Fe resupply after 4 days of Fe deficiency was similar to control plants that had not been subjected to Fe deficiency. Resupply of Fe to the 8 days deficient roots leads to a transcriptional response more closer to the Fe deficient conditions.

Our analysis indicated that 2282 and 1110 genes were significantly up- and down- regulated (p<0.05; LFC>1), respectively, after 4 days of Fe deficiency (Figure 2A, Table S1). Whereas at 8 days of Fe deficiency, 3085 and 2459 genes were significantly up- and down- regulated, respectively (Table S1). Thus, there is an increasing number of DEGs after 8 days of Fe deficiency. In total, 2006 (59.29%) DEGs at 4 days were co-regulated between these time points, with only 150 genes showing opposite expression patterns (Figure 2B, Table S2).

**Figure 2:**
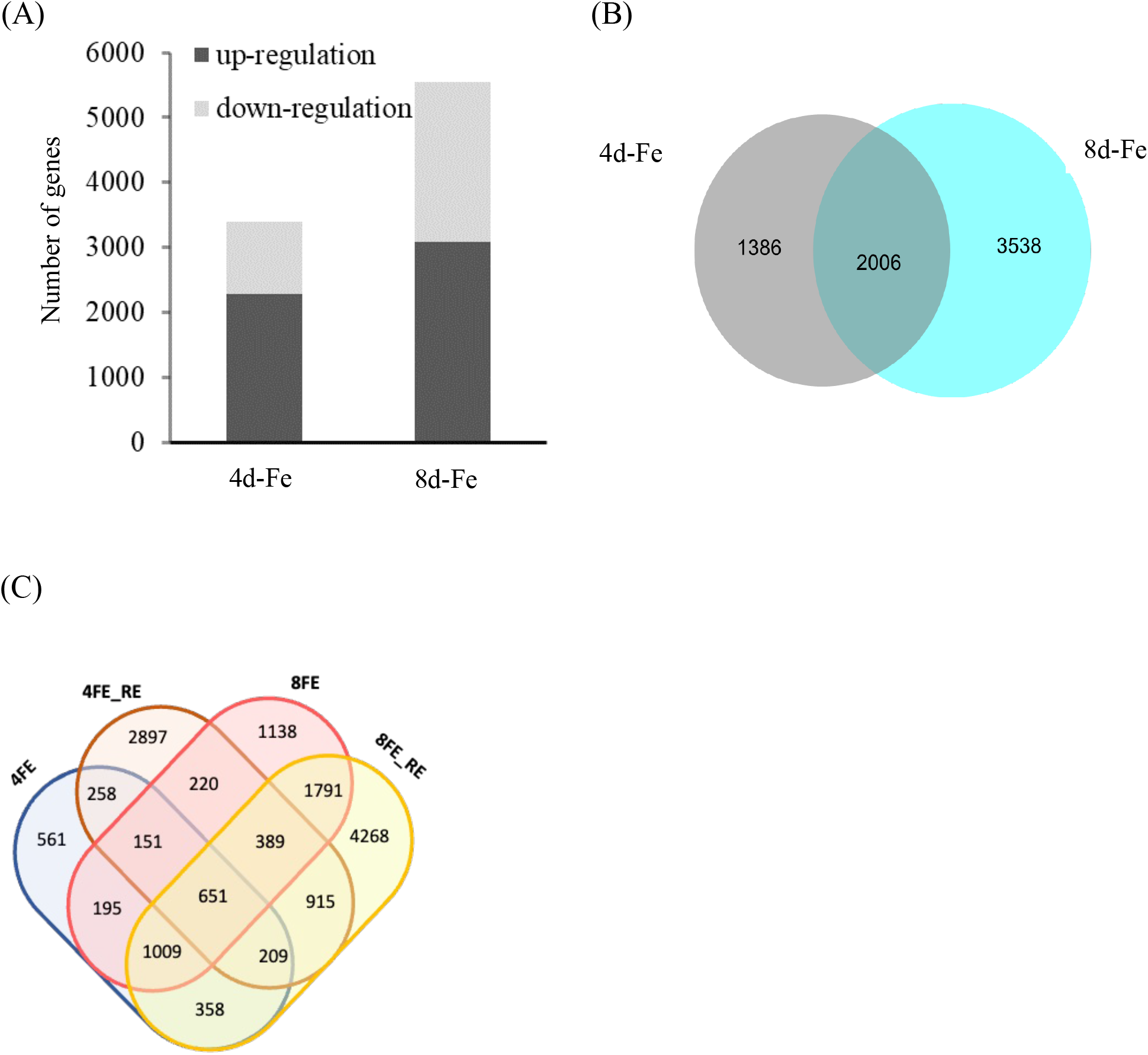
Expression of the identified differentially expressed genes. (A) Total Number of DEGs under the Fe deficiency conditions, and resupply condition. (B) Number of genes those were commonly regulated at 4 and 8 days of Fe deficiency. (C) Venn diagram representing the number of genes for each of the timepoint dataset. The number indicates the genes DE under the mentioned condition.

### 3.2 Genome-wide transcriptional analysis reveals common and distinct clusters of DEGs

To further study the common and distinct responses at the early time points with those observed previously after 20 days of Fe deficiency, cluster analysis of DEG expression was performed. Our analysis revealed eleven different clusters, arranged 1 - 11 by decreasing number of co-expressed genes (Supplementary Figure S3). Some of the clusters contained co-regulated genes largely specific for the 4- and 8 day time points (Cluster 1, 2, 3 and 5), while others contained DEGs in at least two-time points (Cluster 4, 6 to 11). Genes showing upregulation at all three time-points were categorised as Cluster 1, with three sub-clusters consisting of distinct genes with upregulation at each of the 4, 8 and 20 day time points (Figure 3A and Supplementary Figure S3). Within cluster 1, sub-clusters 1a and 1b represents the highest number of genes that are differentially expressed. These include genes encoding for membrane transporters and sulphur-related metabolic processes. Sub-cluster 1c is primarily represented by genes encoding for organic acids and secondary metabolic processes. Interestingly, genes encoding for glutathione-S-transferases were highly expressed in Cluster 1 across all the time points. Clusters 5, 2 and 3 encompass mainly genes which are suppressed after 4, 8 and 20 days of Fe deficiency, respectively (Supplementary Figure S3). Cluster 6 includes genes that are highly induced at 8 days, also including a subcluster of 252 genes that show opposing expression patterns at 8- and 20-days. Comparison with the previously reported RNAseq data from root samples after 20 days of Fe deficiency revealed distinct responses compared to the early time points.

**Figure 3:**
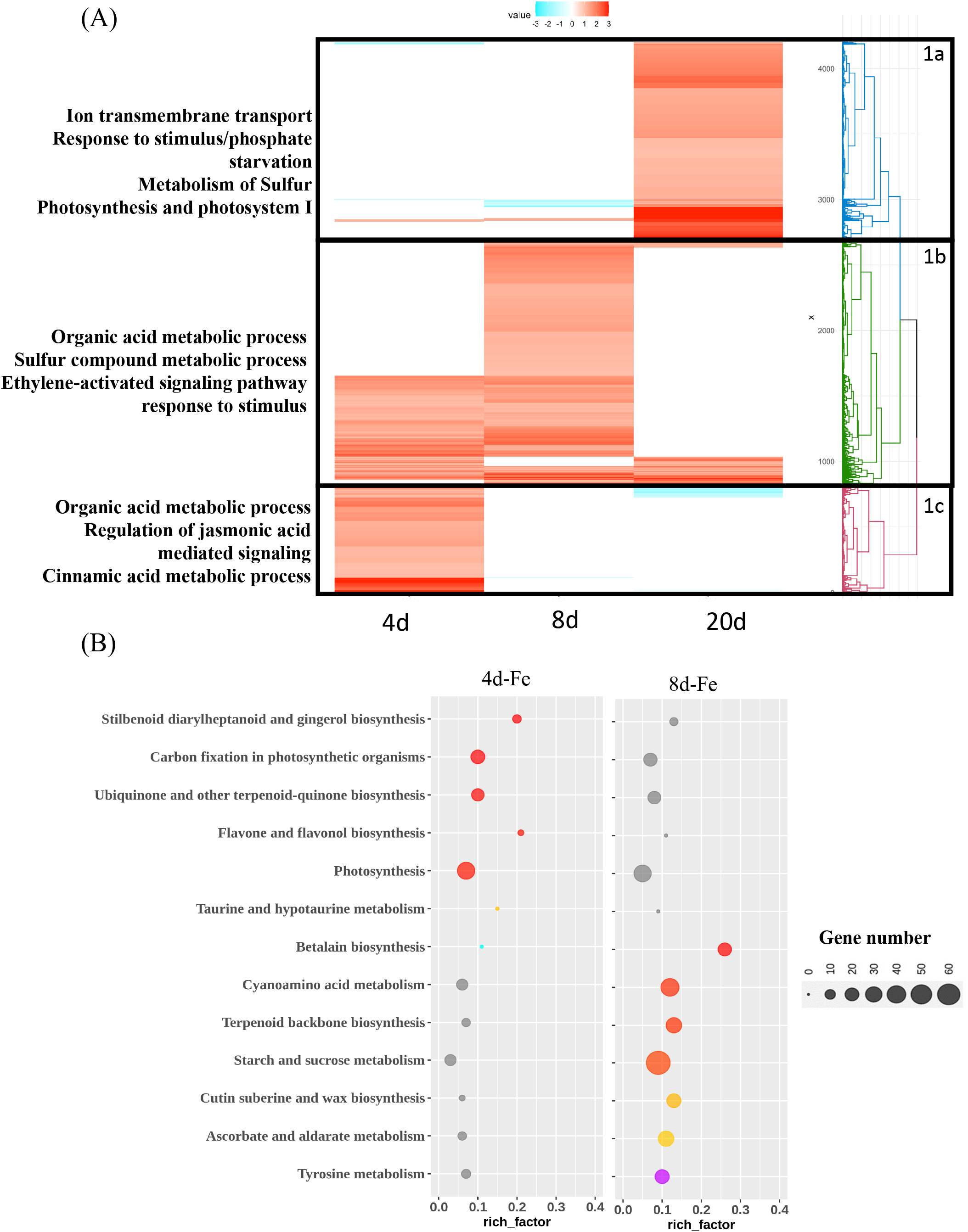
Cluster analysis of DE genes showing distinct time point dependent expression. The total number of genes differentially expressed in roots under after the time points are mentioned while the differentially expressed genes at each time-points show Clustering of the DE transcript expression profiles. Expression profiles (as *Z* scores) of all differentially expressed genes under Fe deficiency condition for each of the time points. (B) KEGG enrichment of the differentially expressed genes with opposing expression patterns during 4 d (4d-Fe) and 8 d (8d-Fe) of Fe deficiency x-axis depicts the rich-factor, i.e., the ratio of perturbed genes in a pathway with respect to the total number of genes involved in the respective pathway, y-axis represents the enriched pathway names, bubble sizes depict the number of genes altered in respective pathways, and increasing intensity of blue color represents increasing significance (decreasing q-value). (Only unique pathway enrichment at time-points represented here; all enriched pathways represented in Supplementary Figure S4).

To understand the characteristic changes in the general and specific functional categories corresponding to early time points of Fe deficiency, we identified significantly over- represented GO terms compared to the controls. This led in identifying 20 GO terms belonging to cellular components, molecular function and biological processes. The early time point shows a higher contribution from functional categories like membrane protein complex, oxidoreductase complex, photosynthetic membrane, envelope, nutrient reservoir activity and response to abiotic stimulus (Supplementary Figure S4A). In contrast, symptomatic time point (8 days) shows a higher number of genes for the GO terms such as cell periphery, apoplast and cellular component organization/biogenesis ((Supplementary Figure S4B).. Similarly, pathway enrichment analysis suggests almost similar major categories at both time points except for phenyl-propanoid pathway, biosynthesis of secondary metabolites and steroid biosynthesis which were highly enriched at 8 days post post-Fe deficiency (Supplementary Figure S5). Pathway enrichment analysis compared to control suggest significant abundance of genes involved in carbon fixation, secondary metabolism and photosynthesis for the 4-day time point (Figure 3B). Specifically, several niche secondary metabolic pathway genes (such as cyanoamino acid, terpenoids biosynthesis, tyrosine metabolism and starch/sucrose metabolism also highly enriched (Figure 3B). For example, genes involved in the biosynthesis of metabolites were highly enriched at the 8 days of Fe deficient roots. Overall, our data indicated multiple functional clusters those showing the cascading effect of gene expression during short-term, symptomatic and long-term Fe deficiency conditions.

### 3.3 Dynamic changes in the gene expression response under Fe resupply

When the Fe-deficient seedlings were subjected to a resupply of Fe for an additional two days, a dramatic transcriptomic rearrangement was apparent. Upon analysis with the respective resupply time points, a total of 1269 (37.41%) and 3840 (69.27%) common DEGs were noted for the 4- and 8-days of Fe deficiency (Figure 2B and C, Table S3). This suggest that a large set of genes were co-regulated at 4 days and 8 days Fe deficiency and only a subset was differentially expressed upon Fe resupply (Figure 4A). We specifically searched for genes that showed reversal of the transcriptional response upon Fe resupply. In the Fe resupply sample after 4 days of Fe deficiency, 625 genes showed reversed expression, while after 8 days of Fe deficiency only 115 genes showed an opposite direction of expression (Supplementary Table S3 and Table 1). This suggest that a longer duration of Fe deficiency could lead to delayed adaptation to the changes in the rhizosphere. We also investigated the expression of already known Fe responsive genes including membrane transporters for their ability to respond to Fe status in roots. Some of the important genes responding to the resupply conditions include DMA biosynthesis pathway genes such as NAS, putative PS exporters annotated as Zinc Induced Facilitator-Like (ZIFL) proteins and multiple YSL membrane transporters (Table S3 and Figure 4B). This specific Fe sensitive response was represented with a higher number of genes at 8 days post Fe resupply (Table S3 and Figure 4). Interestingly, IRT encoding genes (TraesCS4A02G025400, TraesCS7A02G340000, and TraesCS4D02G277100) were up- regulated at 8 days of Fe deficiency and were subsequently downregulated after Fe resupply (Table S3). This suggest that IRT genes are involved during Fe deficiency and its maintenance. Gene enrichment analysis for the data set of Fe resupply after 4 days of deficiency indicates enrichment of genes involved in phenylpropanoid metabolism, plant hormone signalling and signalling pathways. The set of DE genes for resupply, after 8 days of deficiency showed enrichment of genes involved in photosynthesis (Figure 4B).

**Table 1:**
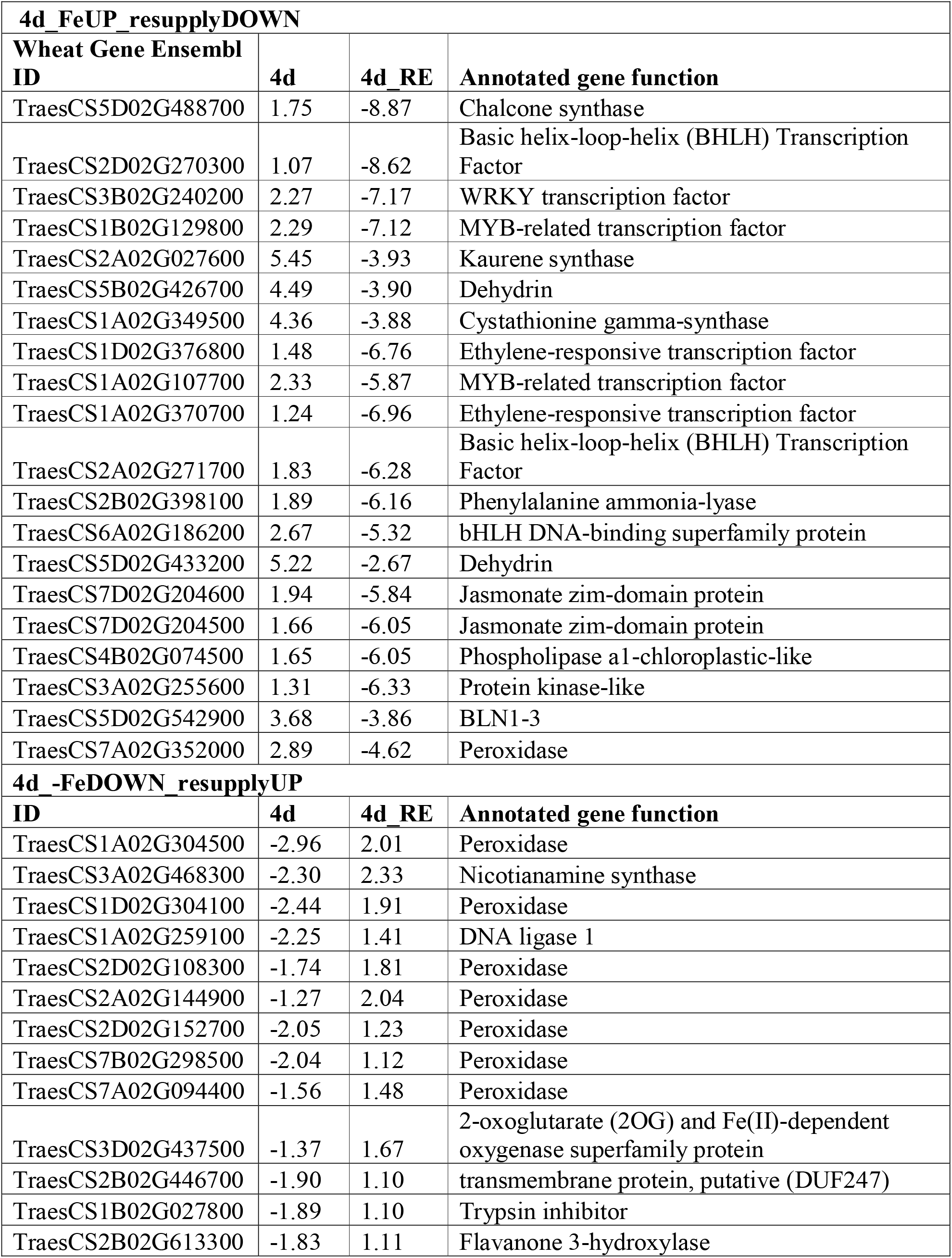
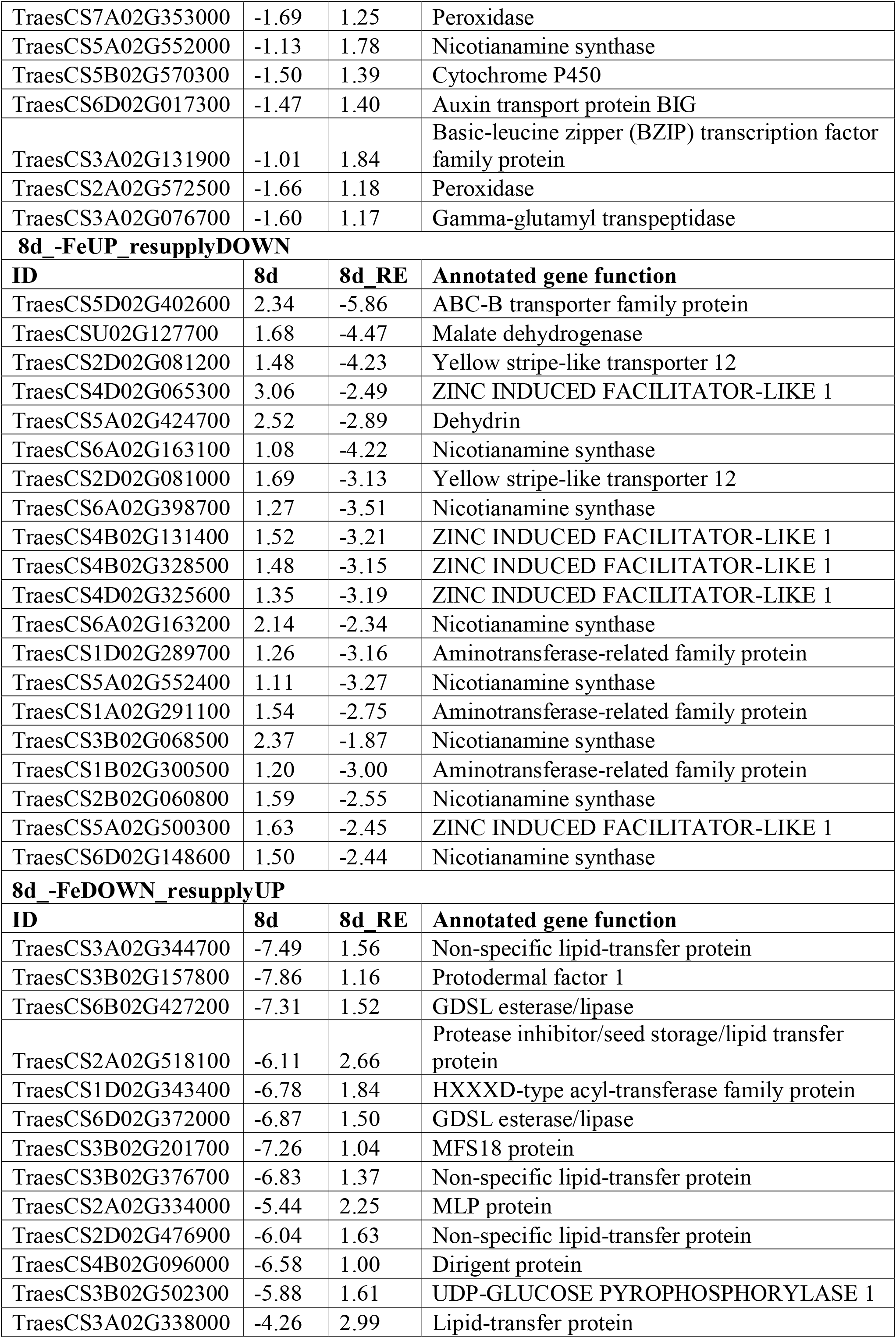
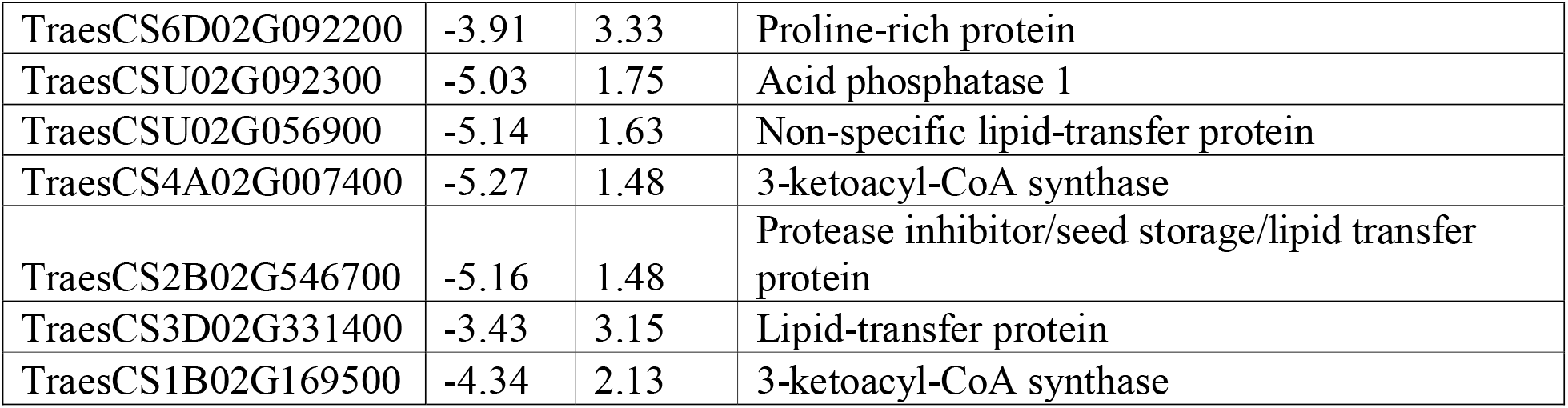
DE genes that showed an opposite expression pattern comparing Fe deficiency (at 4 days and 8 days) and subsequent Fe resupply for 2 days. For each category, only the top 20 genes are shown.

| **4d_FeUP_resupplyDOWN** |  |  |  |
| --- | --- | --- | --- |
| **Wheat Gene Ensembl ID** | **4d** | **4d_RE** | **Annotated gene function** |
| TraesCS5D02G488700 | 1.75 | -8.87 | Chalcone synthase |
| TraesCS2D02G270300 | 1.07 | -8.62 | Basic helix-loop-helix (BHLH) Transcription Factor |
| TraesCS3B02G240200 | 2.27 | -7.17 | WRKY transcription factor |
| TraesCS1B02G129800 | 2.29 | -7.12 | MYB-related transcription factor |
| TraesCS2A02G027600 | 5.45 | -3.93 | Kaurene synthase |
| TraesCS5B02G426700 | 4.49 | -3.90 | Dehydrin |
| TraesCS1A02G349500 | 4.36 | -3.88 | Cystathionine gamma-synthase |
| TraesCS1D02G376800 | 1.48 | -6.76 | Ethylene-responsive transcription factor |
| TraesCS1A02G107700 | 2.33 | -5.87 | MYB-related transcription factor |
| TraesCS1A02G370700 | 1.24 | -6.96 | Ethylene-responsive transcription factor |
| TraesCS2A02G271700 | 1.83 | -6.28 | Basic helix-loop-helix (BHLH) Transcription Factor |
| TraesCS2B02G398100 | 1.89 | -6.16 | Phenylalanine ammonia-lyase |
| TraesCS6A02G186200 | 2.67 | -5.32 | bHLH DNA-binding superfamily protein |
| TraesCS5D02G433200 | 5.22 | -2.67 | Dehydrin |
| TraesCS7D02G204600 | 1.94 | -5.84 | Jasmonate zim-domain protein |
| TraesCS7D02G204500 | 1.66 | -6.05 | Jasmonate zim-domain protein |
| TraesCS4B02G074500 | 1.65 | -6.05 | Phospholipase a1-chloroplastic-like |
| TraesCS3A02G255600 | 1.31 | -6.33 | Protein kinase-like |
| TraesCS5D02G542900 | 3.68 | -3.86 | BLN1-3 |
| TraesCS7A02G352000 | 2.89 | -4.62 | Peroxidase |
| **4d_-FeDOWN_resupplyUP** |  |  |  |
| **ID** | **4d** | **4d_RE** | **Annotated gene function** |
| TraesCS1A02G304500 | -2.96 | 2.01 | Peroxidase |
| TraesCS3A02G468300 | -2.30 | 2.33 | Nicotianamine synthase |
| TraesCS1D02G304100 | -2.44 | 1.91 | Peroxidase |
| TraesCS1A02G259100 | -2.25 | 1.41 | DNA ligase 1 |
| TraesCS2D02G108300 | -1.74 | 1.81 | Peroxidase |
| TraesCS2A02G144900 | -1.27 | 2.04 | Peroxidase |
| TraesCS2D02G152700 | -2.05 | 1.23 | Peroxidase |
| TraesCS7B02G298500 | -2.04 | 1.12 | Peroxidase |
| TraesCS7A02G094400 | -1.56 | 1.48 | Peroxidase |
| TraesCS3D02G437500 | -1.37 | 1.67 | 2-oxoglutarate (2OG) and Fe(II)-dependent oxygenase superfamily protein |
| TraesCS2B02G446700 | -1.90 | 1.10 | transmembrane protein, putative (DUF247) |
| TraesCS1B02G027800 | -1.89 | 1.10 | Trypsin inhibitor |
| TraesCS2B02G613300 | -1.83 | 1.11 | Flavanone 3-hydroxylase |
| TraesCS7A02G353000 | -1.69 | 1.25 | Peroxidase |
| TraesCS5A02G552000 | -1.13 | 1.78 | Nicotianamine synthase |
| TraesCS5B02G570300 | -1.50 | 1.39 | Cytochrome P450 |
| TraesCS6D02G017300 | -1.47 | 1.40 | Auxin transport protein BIG |
| TraesCS3A02G131900 | -1.01 | 1.84 | Basic-leucine zipper (BZIP) transcription factor family protein |
| TraesCS2A02G572500 | -1.66 | 1.18 | Peroxidase |
| TraesCS3A02G076700 | -1.60 | 1.17 | Gamma-glutamyl transpeptidase |
| **8d_-FeUP_resupplyDOWN** |  |  |  |
| **ID** | **8d** | **8d_RE** | **Annotated gene function** |
| TraesCS5D02G402600 | 2.34 | -5.86 | ABC-B transporter family protein |
| TraesCSU02G127700 | 1.68 | -4.47 | Malate dehydrogenase |
| TraesCS2D02G081200 | 1.48 | -4.23 | Yellow stripe-like transporter 12 |
| TraesCS4D02G065300 | 3.06 | -2.49 | ZINC INDUCED FACILITATOR-LIKE 1 |
| TraesCS5A02G424700 | 2.52 | -2.89 | Dehydrin |
| TraesCS6A02G163100 | 1.08 | -4.22 | Nicotianamine synthase |
| TraesCS2D02G081000 | 1.69 | -3.13 | Yellow stripe-like transporter 12 |
| TraesCS6A02G398700 | 1.27 | -3.51 | Nicotianamine synthase |
| TraesCS4B02G131400 | 1.52 | -3.21 | ZINC INDUCED FACILITATOR-LIKE 1 |
| TraesCS4B02G328500 | 1.48 | -3.15 | ZINC INDUCED FACILITATOR-LIKE 1 |
| TraesCS4D02G325600 | 1.35 | -3.19 | ZINC INDUCED FACILITATOR-LIKE 1 |
| TraesCS6A02G163200 | 2.14 | -2.34 | Nicotianamine synthase |
| TraesCS1D02G289700 | 1.26 | -3.16 | Aminotransferase-related family protein |
| TraesCS5A02G552400 | 1.11 | -3.27 | Nicotianamine synthase |
| TraesCS1A02G291100 | 1.54 | -2.75 | Aminotransferase-related family protein |
| TraesCS3B02G068500 | 2.37 | -1.87 | Nicotianamine synthase |
| TraesCS1B02G300500 | 1.20 | -3.00 | Aminotransferase-related family protein |
| TraesCS2B02G060800 | 1.59 | -2.55 | Nicotianamine synthase |
| TraesCS5A02G500300 | 1.63 | -2.45 | ZINC INDUCED FACILITATOR-LIKE 1 |
| TraesCS6D02G148600 | 1.50 | -2.44 | Nicotianamine synthase |
| **8d_-FeDOWN_resupplyUP** |  |  |  |
| **ID** | **8d** | **8d_RE** | **Annotated gene function** |
| TraesCS3A02G344700 | -7.49 | 1.56 | Non-specific lipid-transfer protein |
| TraesCS3B02G157800 | -7.86 | 1.16 | Protodermal factor 1 |
| TraesCS6B02G427200 | -7.31 | 1.52 | GDSL esterase/lipase |
| TraesCS2A02G518100 | -6.11 | 2.66 | Protease inhibitor/seed storage/lipid transfer protein |
| TraesCS1D02G343400 | -6.78 | 1.84 | HXXXD-type acyl-transferase family protein |
| TraesCS6D02G372000 | -6.87 | 1.50 | GDSL esterase/lipase |
| TraesCS3B02G201700 | -7.26 | 1.04 | MFS18 protein |
| TraesCS3B02G376700 | -6.83 | 1.37 | Non-specific lipid-transfer protein |
| TraesCS2A02G334000 | -5.44 | 2.25 | MLP protein |
| TraesCS2D02G476900 | -6.04 | 1.63 | Non-specific lipid-transfer protein |
| TraesCS4B02G096000 | -6.58 | 1.00 | Dirigent protein |
| TraesCS3B02G502300 | -5.88 | 1.61 | UDP-GLUCOSE PYROPHOSPHORYLASE 1 |
| TraesCS3A02G338000 | -4.26 | 2.99 | Lipid-transfer protein |
| TraesCS6D02G092200 | -3.91 | 3.33 | Proline-rich protein |
| TraesCSU02G092300 | -5.03 | 1.75 | Acid phosphatase 1 |
| TraesCSU02G056900 | -5.14 | 1.63 | Non-specific lipid-transfer protein |
| TraesCS4A02G007400 | -5.27 | 1.48 | 3-ketoacyl-CoA synthase |
| TraesCS2B02G546700 | -5.16 | 1.48 | Protease inhibitor/seed storage/lipid transfer protein |
| TraesCS3D02G331400 | -3.43 | 3.15 | Lipid-transfer protein |
| TraesCS1B02G169500 | -4.34 | 2.13 | 3-ketoacyl-CoA synthase |

**Figure 4:**
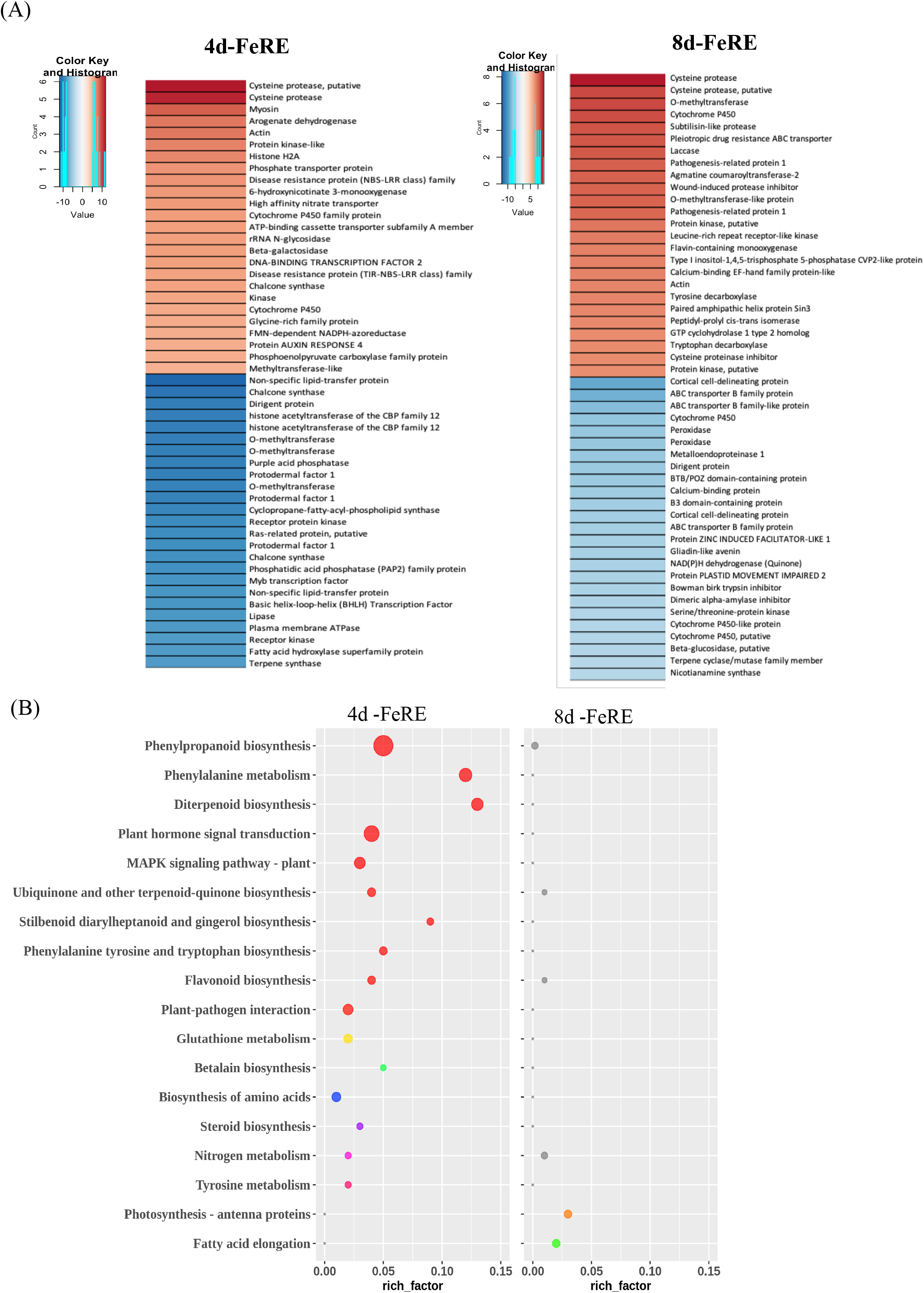
Cluster analysis of DE genes showing distinct time point dependent expression. (A) Heatmap analysis of top 50 up- and down-regulated genes after resupply of Fe to the at 4d and 8d deficient wheat roots. The total number of genes differentially expressed in roots under after the time points are mentioned in Table S3. (B) KEGG enrichment of the differentially expressed genes with opposing expression patterns after resupply of Fe post d (4d-FeRE) and 8 d (8d-FeRE) of its deficiency. The x-axis depicts the rich-factor, i.e., the ratio of perturbed genes in a pathway with respect to the total number of genes involved in the respective pathway, y-axis represents the enriched pathway names, bubble sizes depict the number of genes altered in respective pathways, and increasing intensity of blue color represents increasing significance (decreasing q-value). (Only unique pathway enrichment at time-points represented here; all enriched pathways represented in Supplementary Figure S4).

MFS family genes along with the membrane transporters are a good candidate to sense the exogenous levels of Fe in the rhizosphere. Therefore, to identify the responses specific to the resupply, we extended our study to investigate the expression of the MFS and transporter gene families. Multiple genes belonging to the MFS transcripts encoding for nitrate, phosphate, sugar and oligo-peptide transporters that showed Fe status status-dependent expression response at 4 and 8 days of deficiency (Table S3 and Supplementary Figure S6A). YSL genes, were earlier characterized for the response to Fe deficiency (Kaur et al., 2019; Kumar et al., 2019) yet, their unique response to resupply was intriguing. Interestingly, multiple YSL transporters show reversed gene expression specifically during Fe resupply at both the time points (Figure 5A). Our analysis suggests that *YS1A, YSL12, YSL13, YSL19* and *YSL20* show expression response co-related with the Fe concentrations in the rhizosphere at either of the time points (Figure 5B).

**Figure 5:**
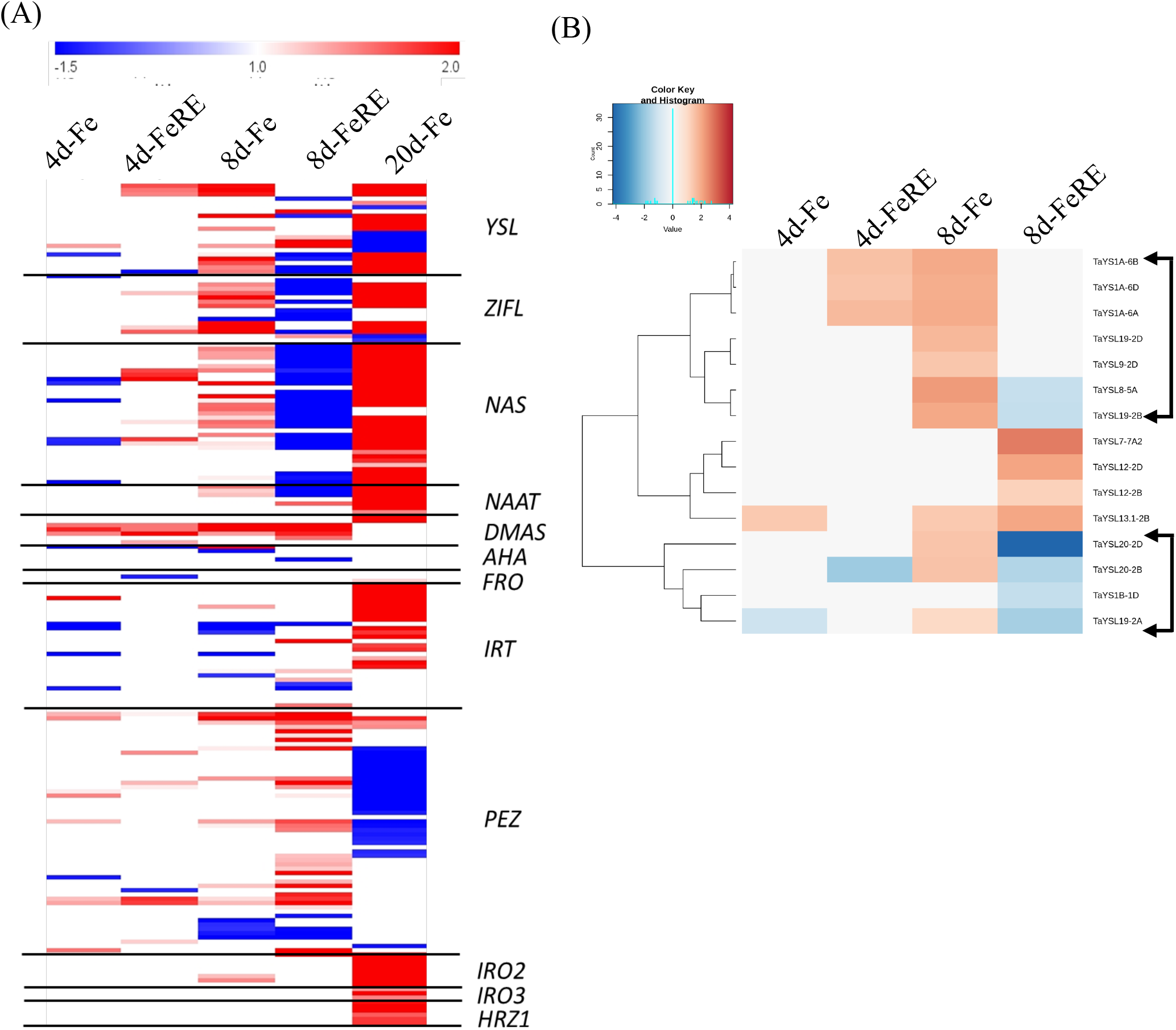
Analysis of FSR and YSL transporters genes. **(A)** Heatmap analysis the Fe-responsive genes and their abundance in the wheat roots grown under different Fe-status. Changes in the response of the Fe-deficiency were noted with a p-values of <0.05 were selected for analysis. (B) The heatmap analysis of the wheat YSL transporters showing differential regulation under the Fe-deficiency conditions.

To identify the unique regulatory signatures for the time points, an analysis of the expression levels of the TFs was performed. As expected, differential induction of multiple TFs family genes of AP2-EREBP, WRKY, bHLH and MYB was observed (Supplementary Figure S7). The highest number of TFs encoding genes (371) were differentially regulated after 8 days of Fe deficiency. In all, 723 TFs were found to be differentially regulated in response to Fe deficiency at either of the three time-points (Table 2). Distinctively, the highest number of TFs (1192 genes) were perturbed in resupply condition (Table S4). Interestingly, only 14 TFs are coregulated at 4, 8 and 20 days of Fe deficiency (Table S4b). Specifically, 72, 207 and 264 TFs were DE exclusive for Fe deficiency at 4, 8 and 20 days, respectively. Our heatmap analysis suggests that a large number of TFs show Fe-resupply sensitive responses that occurs primarily at the early stage of Fe sensing (Supplementary Figure S7). Expression data also reveal that during 4 days of Fe deficiency and its resupply treatment, gene expression of 62 TFs was were either reverted to or down-regulated to adapt the Fe-flux. In contrast, this phenomenon of reversal gene expression was not visible at the 8 days of Fe deficiency compared to the resupply condition. Interestingly, Arabidopsis Fer-like iron deficiency-induced transcription factor (FIT) homolog in wheat TraesCS2A02G271700 was downregulated upon Fe resupply (Table 1). Overall, our data points towards distinct core molecular respond stringently to the Fe root status.

**Table 2:** Number of TF gene families those were differentially regulated [up-regulated (up) or down-regulated (down)] during the two time points (4 and 8 days) of Fe deficiency (–Fe) and post Fe-resupply.

| TFs categories | Number of genes |  |  |  |
| --- | --- | --- | --- | --- |
|  | Common DEGs in –Fe (4D or 8 D) |  | Common DEGs in Fe_RE (4D or 8 D) |  |
|  | Up | Down | Up | Down |
| bHLH | 23 | 23 | 34 | 46 |
| WRKY | 57 | 1 | 32 | 28 |
| MYB | 47 | 12 | 59 | 16 |
| AP2-EREBP | 56 | 14 | 52 | 52 |
| C2H2 | 13 | 6 | 39 | 18 |
| NAC | 11 | 5 | 17 | 20 |
| MADS | 7 | 2 | 12 | 5 |
| GRAS | 5 | 0 | 10 | 6 |
| PHOR1 | 20 | 0 | 19 | 1 |

### 3.4 Fe status affects the genome induction bias in hexaploid wheat roots

We expanded our study to perform a comparative analysis of induction bias from the datasets generated at early time points of Fe deficiency, resupply and late phase (Kaur et al., 2019). The triplets contributing towards altered transcriptome expression can be divided into (i) 3up/3down: the ones showing similar altered expression across all the homoeolog triplets, (ii) 1up/1down/2up/2down: triads showing biased altered expression across the triplet genes, and (iii) Opposite: homoeolog genes within a triad showing opposing expression patterns. Our analyses suggest that a major proportion of the homeolog triplets show no differential expression (No DE; Figure 6, Supplementary Table S5). In general, the percentage of differentially expressed triads (DETs) increases with the duration of Fe stress, with 7.78% and 12.91% of the accepted triads showing DE at 4 and 8 days of Fe deficiency, respectively. Among the DETs majority depict induction biasness patterns (80.36% at 4- and 76.42% at 8- days), with ∼50% of the DETs contributing to single homoeolog induction (1bias) and ∼25% showing two homoeologs induction (2bias). In comparison, a lower percentage of triads were induction biased in resupply samples. Only 7.41% of the DETs at 4 days showed the same response upon Fe resupply, which increased to 33.41% at 8 days. Remaining DETs showing dissimilar patterns were analyzed for GO enrichment. Interestingly, ∼22% of the 4 d DETs show same patterns at 8 d also, while for resupply time-points only ∼5% had similar patterns. Compared to early Fe deficiency time-points, the similarity is far lower across early and late time-points (4- and 20 days with 3.65%; 8 d and 20 d with 2.67%). When contribution from the sub-genomes was calculated for the 1bias category, B genome contributed the most (4 days, 35.33% and 8 days, 34.88%), followed closely by A genome (4 days, 34.77% and 8 days, 33.17%) for early time-points while the contribution was higher from the A genome for the late Fe deficiency time-point. The sub-genomic contribution in the 2bias triads show a higher B+D sub genome bias at early time-points, while a higher A+D bias at 20 days of time points (Table S5).

**Figure 6:**
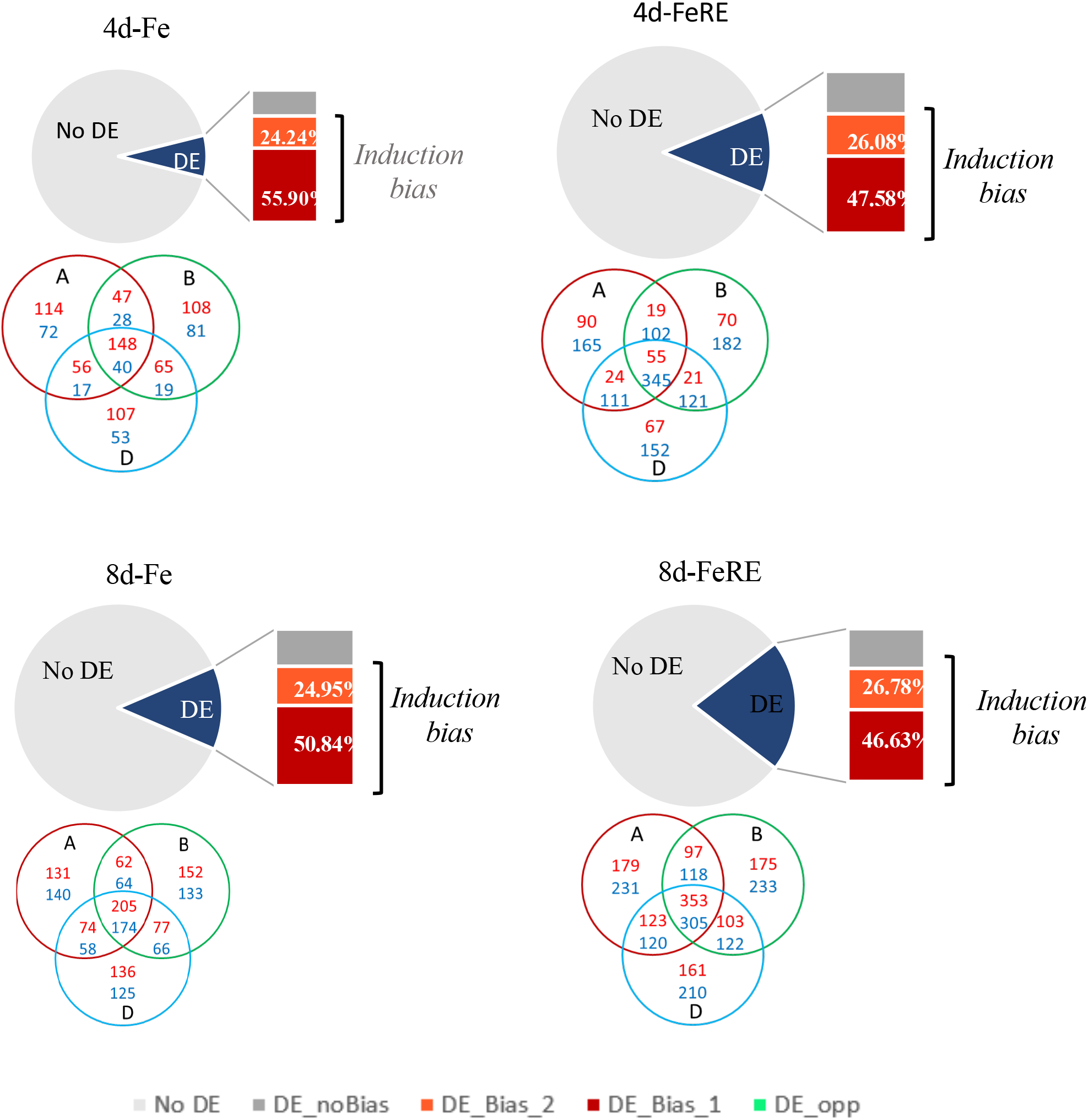
Induction biasness patterns for the Fe-deficiency as well as resupply datasets. Figure summarizes the percentage of accepted triads that were differentially expressed (DE) at 4 d of Fe deficiency (4d-Fe), 4d-Fe followed by 2 d of Fe resupply (4d-FeRE), 8 d of Fe deficiency (8d-Fe), 8d-Fe followed by 2 d of Fe resupply (8d-FeRE). For each of these conditions, the differentially expressed triads (DETs) were categorized as those showing no bias across the homoeologs (DE_noBias), and those showing induction bias (DE_Bias_1: 1 homoeolog showing DE; and DE_Bias_2: 2 homoeologs showing DE). Further, for each of the conditions the contributing genomes for the Induction biased triads are represented through venn diagrams. Red color denotes “A” homoeologs, green: “B”, blue: “D”.

We also came across a few triads with homeologs showing opposite expression induction. Two triads show opposing expression trend at 4 days and 10 triads at 8 days of Fe deficiency, while upon resupply, 2 and 20 triads show opposite induction at 4- and 8-days respectively (Supplementary Table S6). These include trypsin inhibitor and a kinase at 4 days of Fe deficiency; ethylene-responsive TF and cotton fibre fiber expressed domain-containing protein show at resupply treatments of 4 days of Fe deficiency (Supplementary Table S6); aquaporin-1, sugar, heavy metal, peptide, ammonium transporters, glycosyltransferase, protease inhibitor/seed storage encoding genes at 8 days of Fe deficiency; ethylene-responsive and bZIP TF, CASP-like, WAT1-related, carboxypeptidase, globulin genes, chalcone-flavone isomerase, glycosyltransferase, aminotransferase, NAG, peptide and ammonium transporter at 8 d resupply. In totality, we observe increased opposing induction bias for genome at 8 days when compared to 4 days (Table S5 and S6), with molecular functions related to binding, transporter activity and transcription regulation.

Subsequently, we also analysed the relative expression bias within triads that suggested a similar pattern to the induction bias at 4 d of Fe deficiency, with an increase in balanced expression upon Fe resupply. Across the conditions, B homoeologs were predominantly suppressed while D and A homoeologs show higher dominance (Supplementary Figure S8 and Table S7). We also checked if the response to Fe deficiency is limited to certain regions of the genome, and therefore distribution of the DEGs on the hexaploid wheat genome was analysed. Our analysis indicates a higher contribution from chromosomes 2, 3, 5 and 7 while upon resupply, comparatively higher contribution was observed from chromosome 2. It will be interesting to study the location of these bias triads on the chromosomal unit.

### 3.6 Fe deficiency causes a gradual increase in homoeolog triad expression for transporters

Next, chromosomal mapping of all the triads was done on the chromosome. The induction bias transcripts arising from all the conditions show similar prevalence in high/low recombinant regions of the chromosomes (Figure 7). Specially, we observed an increased percentage of biased genes were localized on proximal chromosomal regions (R1/R3) that undergo high recombination, when compared to the percentage of genes that do not show bias (3flat/3up/3down). While a comparatively lower percentage of low recombinant regions (R2/C) genes show induction bias. These observations for expression bias correlation with the genomic compartments of the wheat chromosomes was in accordance with the previous observation (Ramírez-González et al., 2018). We further went on to perform the comparative induction bias of TFs (1761 triads) and transporters (94d triads) those falling in the triads. Our analysis suggests that the percentage bias was significantly similar for the TFs. This biasness was not consistent for the category of genes encoding for transporters. We observed an enhanced biasness of the triads with the time period of Fe deficiency stress suggesting the compounding temporal effect of the stress on the genome genome-specific expression. This suggests that differential bias expression of the transporters is an important feature to adapt deficiency tolerance in hexaploid wheat.

**Figure 7:**
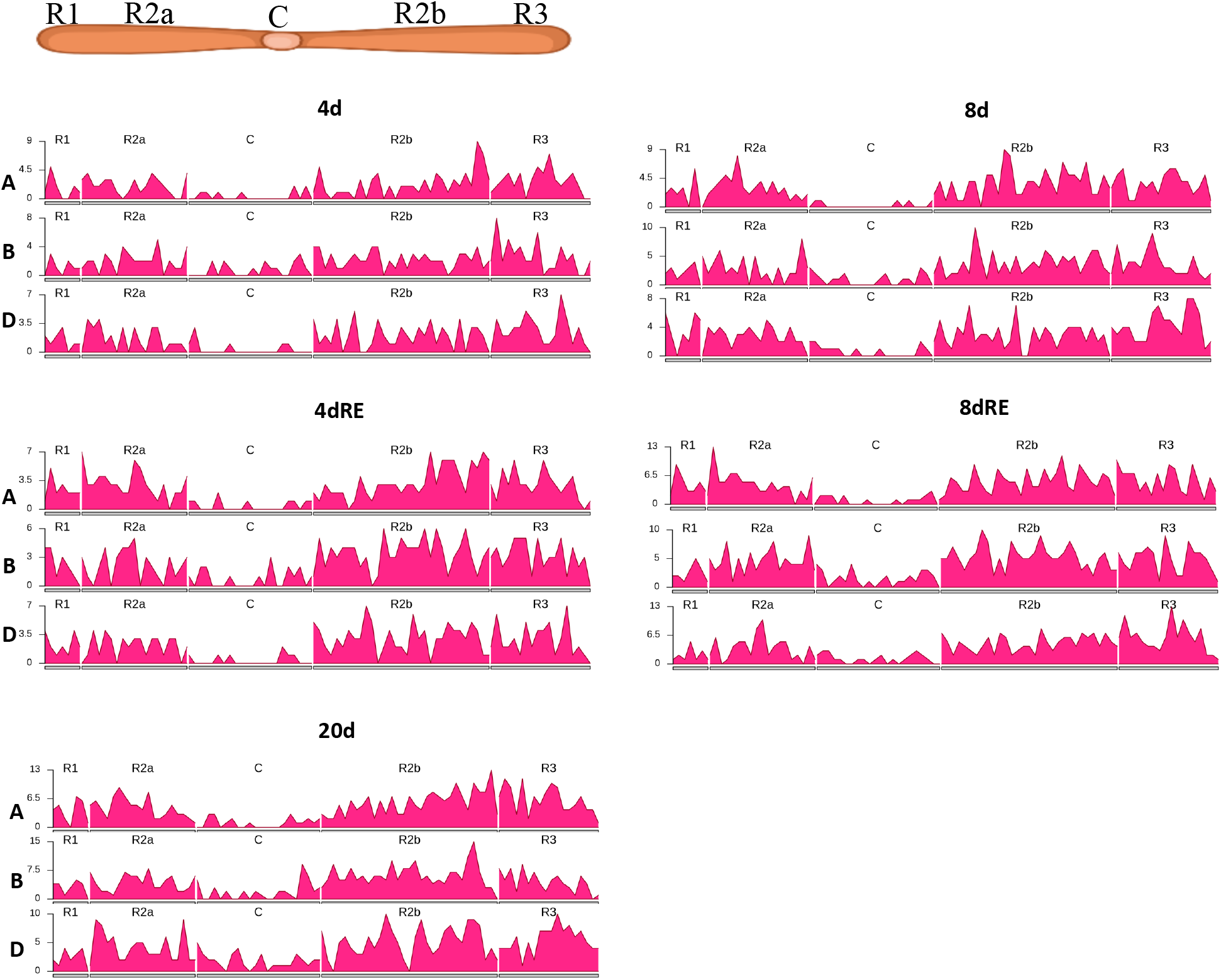
Chromosomal localization of the homoeologs showing biased induction/suppression. The chromosomal localization of differentially expressed gene was determined using wheat Ensemble database (IWGSC RefSeq v1.0, Ensembl release 47). The chromosomal location is associated with distal (R1 and R3) and interstitial and proximal (R2 and C) regions.

## 4. Discussion

This study aimed to identify the distinct changes in the transcriptome of hexaploid wheat during the initial stages of Fe deficiency. Our physiological experiments validated the 4 and 8 days of Fe deficiency as an appropriate time frame to stimulate early response in wheat seedlings. Changing the Fe status during the early stages of plant growth helps in defining the early molecular responses involved in modulating the symptomatic responses. Earlier studies mostly focused on either the late stage of plant development or a longer duration of the Fe deficiency stress (Kaur et al., 2019; Wang et al., 2019). Utilizing RNA-seq analysis could provide an insight in identifying the signals controlling early molecular events and those involved in sensing by resupplying Fe. The present work reveal the distinct response during the time points that manifest the molecular events including the expression of the membrane transporters associated with the nutrient deficiency response.

Our experimental setup showing the growth defects confirming the relay of the root Fe deficiency to the shoot tissue (Figure 1). Transcriptomic analysis revealed the functional categories that include carbon partitioning, remodelling of lipid membranes and Fe mobilization mechanisms those which are characteristic to the early response. Such rearrangement in plants are necessary due to prevent them from the Fe deficiency induced damages. Previously, proteome analysis of rice Fe deficiency response also show changes to Carbon and energy Metabolism (Selby-Pham et al., 2017).

Interestingly, when resupplied with Fe, we observed the down-regulation of major gene families involved in Fe uptake and mobilization mediated through the strategy-II (Figure 5A). In addition, multiple membrane transporters also show down-regulation under Fe resupply condition. Investigating Fe resupply transcriptomics could reveal important sensors and regulators for the establishment of the Fe deficiency responses. In this study many such genes show Fe status dependent expression response. Most importantly, genes primarily involved in DMA biogenesis and its secretion were highly responsive to the Fe status. Such conserved molecular response was noted across the monocots including maize, barley (Bocchini et al., 2015; Zanin et al., 2017). Specifically, we found that genes encoding for wheat TOM1, TOM2 and TOM3 show high expression at 8 days of Fe deficiency with the reversal to the basal expression when resupplied with the Fe (Figure 5A and Table S1 and S3). Thus, resupply of Fe has resulted in minimizing the detrimental effect of its deficiency. In monocots, TOM genes were characterized previously as an essential functional unit for the release of PS in the rhizosphere(Nozoye et al., 2011, 2015; Sharma et al., 2019). We extended this analysis for transporter genes showing sensitivity for Fe resupply. We hypothesized that the down- regulation of genes during resupply condition could represent membrane proteins that could sense the Fe status. This criterion led to the identification of subsets of transporters including IRT1, YSL and metal transporters showing Fe status dependent response in roots at the symptomatic stage. YSL genes belongs to the OPT gene family are known for their role in mobilizing divalent cations including micronutrients such as Fe (Lubkowitz, 2011; Yordem et al., 2011). Earlier, multiple wheat YSL genes were identified and characterized for their expression response to the changing regimes of micronutrients (Kumar et al., 2019). Amongst multiple transporters, YS1A show root Fe status dependent response. Previously, ZmYS1A has been characterized for its binding and transport ability of Fe(III) chelates (Harada et al., 2016). The presence of conserved Fe^3+^-PS binding at the 7^th^ extracellular loop region likewise to HvYS1A suggests that it is able to transport specific forms of Fe (Harada et al., 2007). Previously, TaYS1A was shown to be localized on the cell membrane (Islam et al., 2020). Likewise other plant YS1A homologs, we expect the TaYS1A should also transport Fe(III)-PS. Specifically, *H. vulgare* domains showing high affinity for Fe^3+^-PS compared to other plant species and wheat YS1A showing closest homology to HvYS1A (Banakar et al., 2017; Harada et al., 2007).

In our study, the concerted regulation of the TFs was evident by a large number of DEGs encoding for different classes of TFs. Many studies have suggested the predominance role of bHLH TFs during the Fe deficiency response that remains conserved between plant species (Gao et al., 2019). Multiple studies have demonstrated regulatory-hub role of FIT during Fe homeostasis (Colangelo & Guerinot, 2004; Schwarz & Bauer, 2020). In our analysis FIT induced during early stage of Fe deficiency was down-regulated under resupply condition indicating a possible conserved role of wheat bHLH TF. Multiple rice and Arabidopsis bHLH TF homologs have been identified from hexaploid wheat and they show DE response under Fe deficiency (Kaur et al., 2019; Kumar et al., 2022)

Additionally, in this study, we observed that plant specific WRKY TFs were significantly induced and show a high number of genes with increased duration of Fe deficiency. Majorly, WRKY proteins are known to be involved during different biotic and abiotic stresses (Banerjee & Roychoudhury, 2015). Earlier, AtWRKY46 was shown to be involved specifically during the mobilizing root Fe to shoots by regulating Fe transporters (Yan et al., 2016). The high numbers of genes encoding WRKY TFs in our Fe deficient roots and also during resupply suggest its importance for Fe translocation. Our detailed investigation shows 52.5% of the Fe responsive wheat genes (20 d) having more than one W-box domains in their promoters (Supplementary Table S8). Further studies are required to confirm the functional role of few candidate WRKY TFs regulating specific to the process of Fe flux.

Genome biasness could contribute the transcriptomic landscape in hexaploid wheat during the exposure of biotic and abiotic stresses (Akhunova et al., 2010; Powell et al., 2017). Specifically, during the Fe deficiency, the changes in genome and expression bias offered by A and B genome was significant and forms the basis for further investigation (Kaur et al., 2019; Wang et al., 2020). In this study, we carried out an in-depth analysis for of genome induction and expression bias. A higher number of triads are altered under resupply conditions, a lower percentage of induction biasness is observed under resupply compared to Fe deficient conditions. Interestingly, some of the transporters show opposite induction from the homoeologs. This suggests a definite change in the expression bias in wheat responding the nutrient deficiency such as Fe. Some of the candidate genes encoding for peptide (POT family), heavy metals transporter (HMA) and auxintransporter (WAT1-related protein) show B genome specific induction bias (Table S6). Earlier, these transporters have been reported for their role in Fe homeostasis (Kaur et al., 2019). We also observed that, the response from the specific wheat chromosome (Chromosome 2,3,4 and 7) show homoeologs induction percentage that increases with the duration of the Fe stress. The same analysis was repeated for our previous 20 days of Fe deficiency exposed wheat root dataset (Kaur et al., 2019). Interestingly, A genome genes predominantly contribute towards the 1up/1down induction bias, while A and D homoeologs show a higher contribution for 2up/2down biasness. Our GO term analysis suggests an overall stronger abiotic stress response and nutrient reservoir activity the early deficiency stage, while the symptomatic stage shows changes in structural organization and localization and more prominent deficiency responses that are not reverted upon Fe resupply (Supplementary Figure S4). We observed that, although the same GO categories were induced in Cluster-1 (Figure 3A) yet their different homoeolog derived transcripts were expressed for each time points. Notably, our analysis also reveals the homoeolog triad encoding for transporters show significantly higher biasness when compared to TFs. These results infer the importance of genome bias for modulating the varying expression responses during the stress condition. Such bias in the allopolyploid crop points to the epigenetic changes and are of evolutionary significance that points to the genome interactions especially under stress (Akhunova et al., 2010; Powell et al., 2017; Yoo et al., 2013)

Our result delineates the distinct response during the early phases of Fe-deficiency response establishment (Figure 8). A clear distinct transcriptional at the early and during the onset of phenotypic responses were noted. As compared to the 4 day post Fe deficiency, surmounting expression of membrane transporters, metabolic pathways and TFs was noted after 8 days. Differential transcriptome-reset was observed after the resupply of Fe, with multiple membrane transport showing reversal of gene expression.

**Figure 8:**
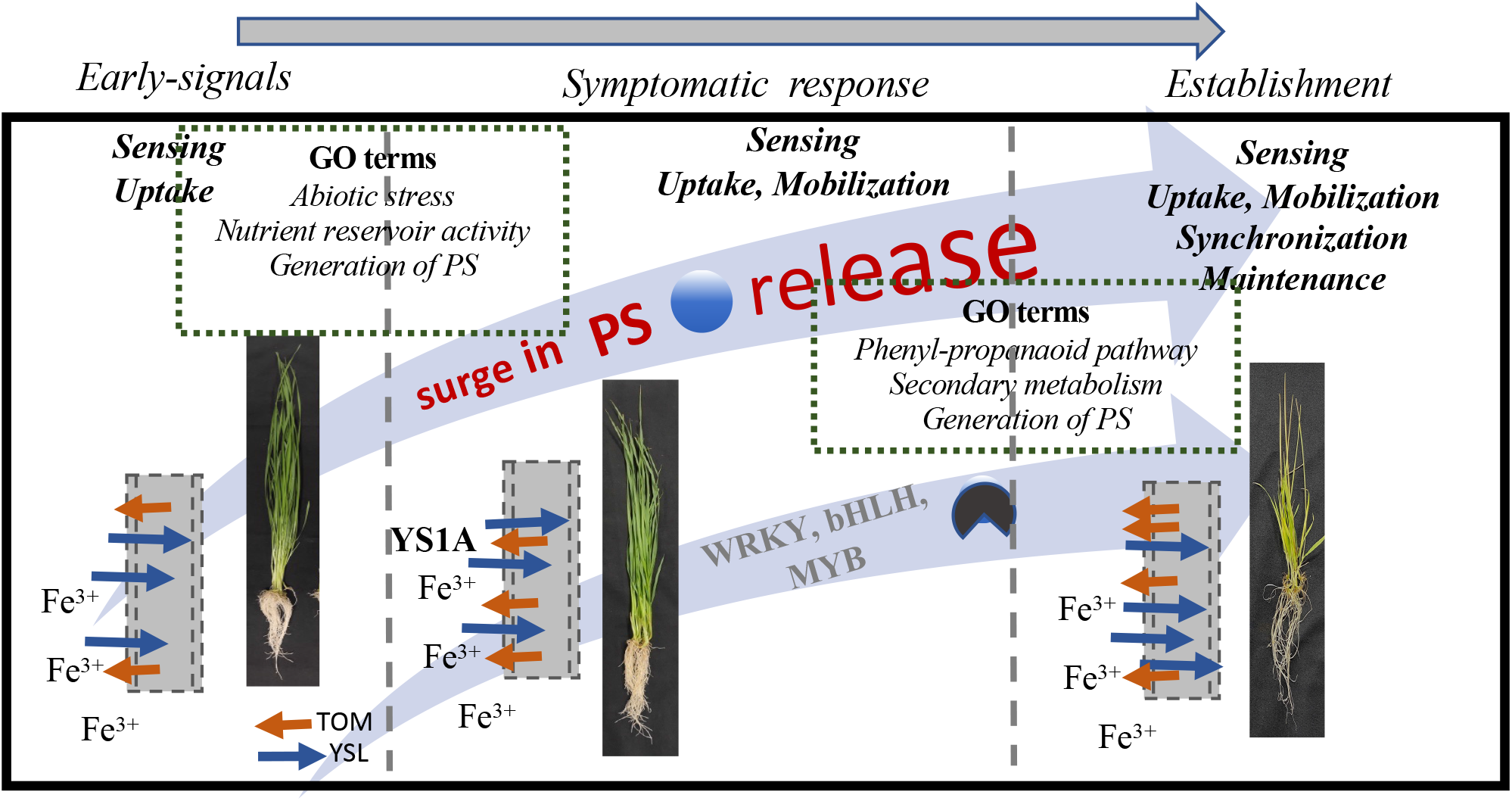
Model representing the event occurring during early, symptomatic and late phase of Fe-deficiency. The early phase (4 d) of Fe-deficiency represents by the GO function such as abiotic stresses, nutrient reservoir activity and generation of PS; the symptomatic response (8 d) is represented by surge in the gene expression involved in phenyl-propanoid biosynthesis pathway and secondary metabolism surge as a Fe-deficiency response. The late phase of deficiency response (20 d) is represented by synchronization of events to mobilize the Fe in hexaploid wheat. The Fe-deficiency response was also accompanied by the surge in the gene expression involved in the biosynthesis and release of PS and gradual expression of the multiple TFs.

## 5. Conclusions

Thus, this study features the distinct elements of the Fe-deficiency response during early signalling stage that show transcript expression burst at the symptomatic stage. Furthermore, the involvement of multiple components to enhance mobilization of Fe by modulating PS release, expression of membrane-transporters that show robust expression with the changing phase of the deficiency. These events help establish and wiring the connections in the hexaploid wheat that could help integrate the functions for surviving the fluctuating levels of Fe. Overall, this which will improve our understanding of the molecular mechanisms underlying early phase of Fe deficiency and its regulation in hexaploid wheat. This will enable researchers to screen for more efficient Fe utilizing efficiency genotypes for the agricultural practises.

## Declaration of Competing Interest

The authors declare that they have no known competing financial interests or personal relationships that could have appeared to influence the work reported in this paper.

### CRediT authorship contribution statemen

AKP, JB and VM conceived and designed the study. VM, RJ, GK and AK carried out the experiments and performed the morphological analysis; GK and VM performed the RNAseq analysis; GK, JB, VM, RJ, and AKP analysed data and wrote the paper. AKP, JB, VM and GK reviewed and edited the manuscript. All authors read and approved the final manuscript.

### Data availability

All the data generated in this study are included in this publishing article and its supplementary information.

## Acknowledgments

The authors thank the Executive Director for facilities and Dr. Sophie Harrington for valuable discussions. This work was supported by NABI-CORE grant to AKP. DBT-eLibrary Consortium (DeLCON) is acknowledged for providing timely support and access to e-resources for this work.

## Legends for Supplementary data

Table S1: List of genes showing differential expression at 4 d (upregulated: Sheet 1, downregulated Sheet 2) and 8 d (upregulated: Sheet 3, downregulated Sheet 4) of Fe deficiency w.r.t. respective control root samples.

Table S2: DEGs showing opposite expression patterns at 4 d and 8 d of Fe deficiency w.r.t. respective controls. Sheet 1 enlists 8 genes induced at 4 d, while repressed at 8 d and Sheet 2 lists 142 genes those are repressed expression at 4d but induced at 8 d of Fe deficiency w.r.t. respective control root samples.

Table S3: DEGs depicting opposite expression patterns upon Fe resupply.

Table S4: List of Transcription Factors those are differentially expressed at different Fe deficient and resupply conditions w.r.t. control root samples.

Table S5: Homoeolog triad *induction* bias patterns in response to different Fe deficiency and resupply conditions w.r.t. respective control samples.

Table S6: Details for homeolog triads with opposing homoeolog induction bias in response to different conditions w.r.t. control.

Table S7: Relative expression values were calculated for the gene triads for the control as well as –Fe datasets revealing D genome to be contributing the most towards overall expression, followed by A and B genomes.

Table S8: Fe responsive DEGs those show present of W-box domains. The DEGs were searched for the presence of the w-box motif (T)TGAC(C/T) in the 2000 bp upstream promoter regions using an in-house perl script.

