## Supplementary Figure S1_S8 for "Asymmetric expression of homoeologous genes in wheat roots modulates the early phase of iron-deficiency signalling"

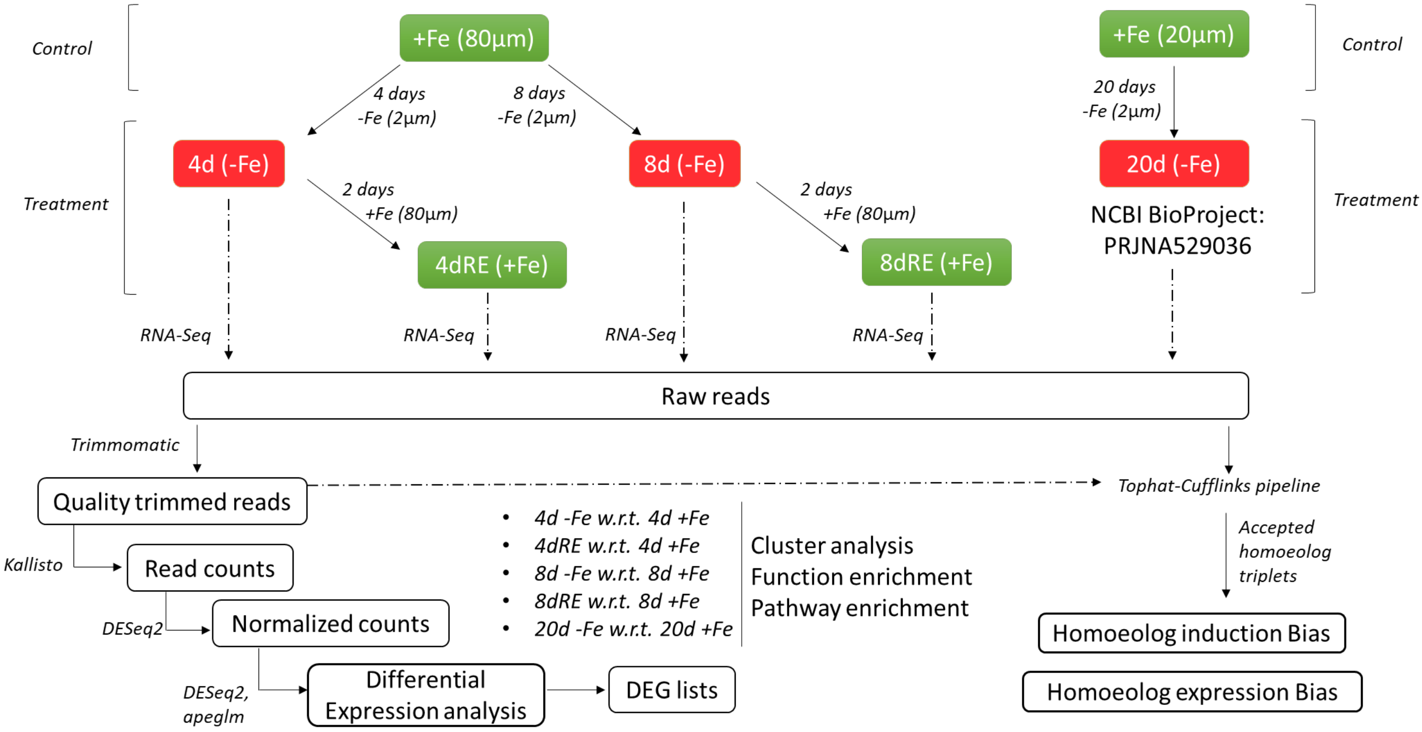


**Figure S1:** Schematic representation of the comparative RNA-Seq analysis across different conditions and time-points.

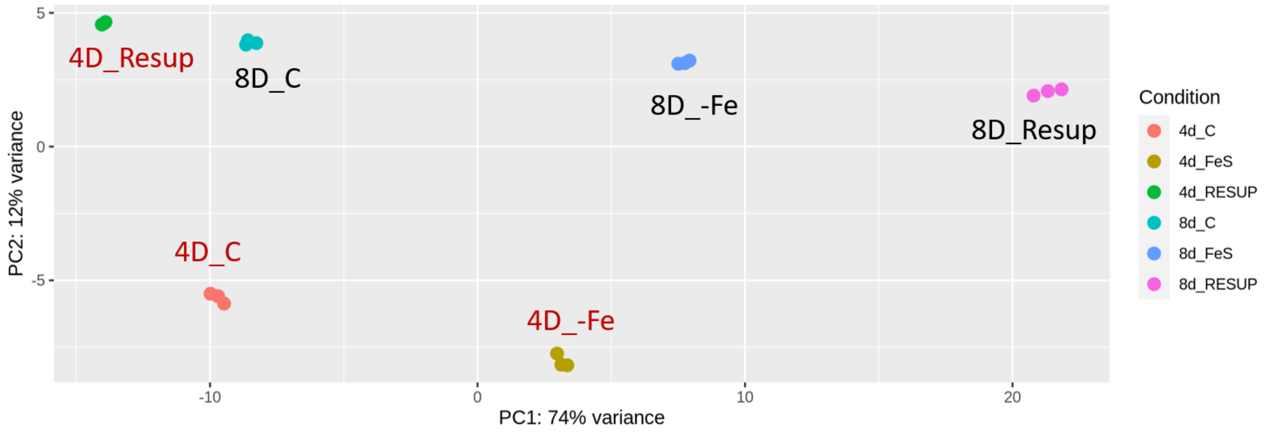


**Figure S2:** Principal Component Analysis for different conditions and time-points. The plot indicates that biological variation for replicates and different samples. 4D_C and 8D_C indicates control root samples; whereas 4D_Fe and 8D_Fe indicates wheat root transcriptome generated in Fe deficiency condition. 4D_Resup and 8D_Resup indicated the resupply of Fe to the respective deficiency treatments for two days.

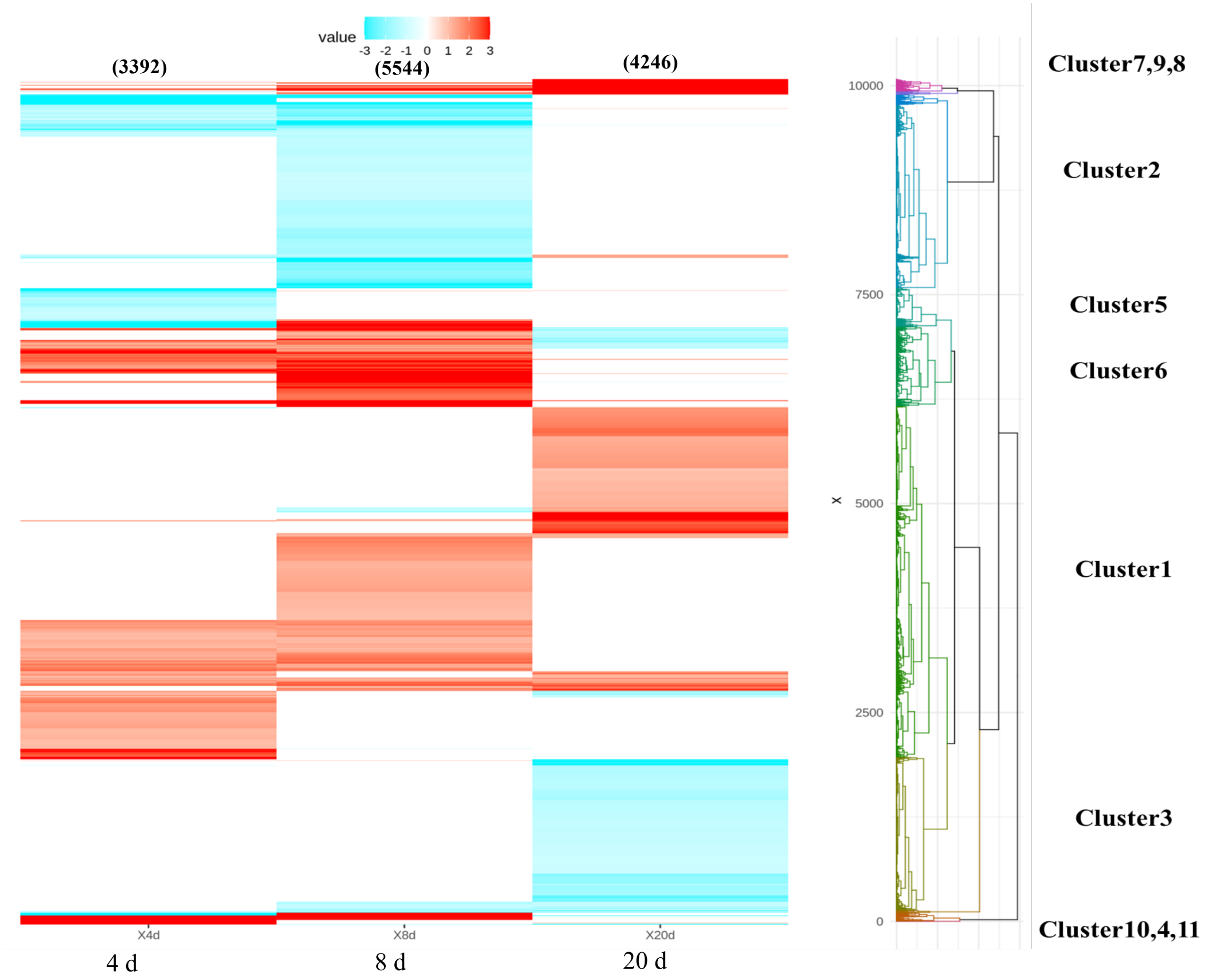


**Figure 3:** Expression profiles of all differentially expressed genes from roots at 4, 8 (this study) and 20 days (Kaur et al., 2019) post Fe deficiency. The total number of genes differentially expressed in roots under after the time points are mentioned while the differentially expressed genes at each time-point Clustering of the DE transcript expression profiles.

(A)

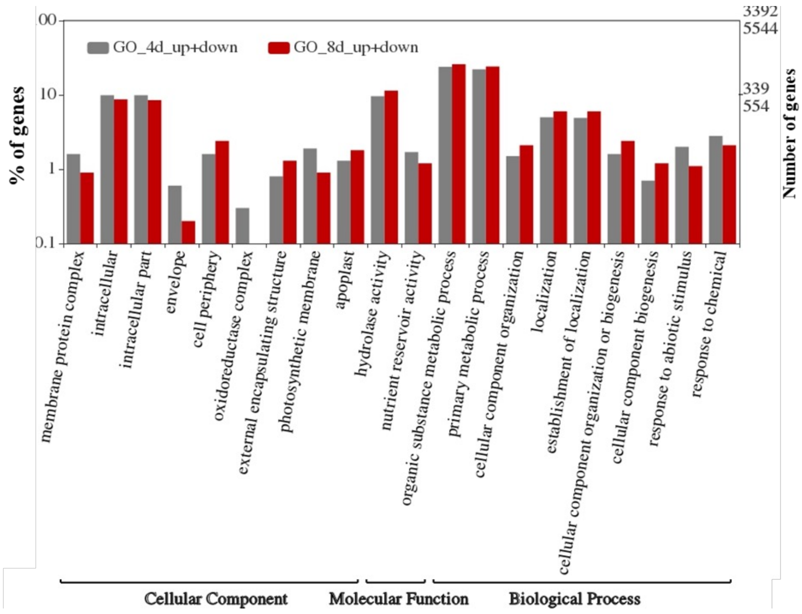


(B)

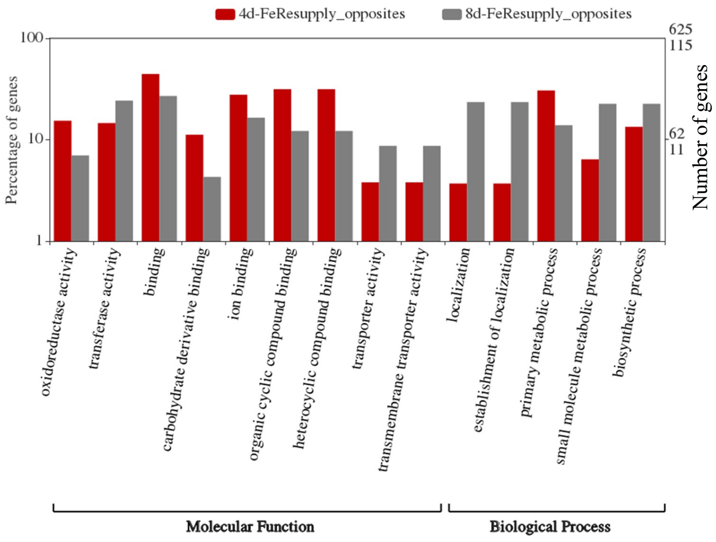


**Figure S4: Functional categorization analysis of the DEGs.** (A) Gene Ontology (GO) categorization of the DEGs during 4 and 8 d of Fe deficiency compared to controls. (B) GO categorization of the DEGs showing opposing expression patterns upon resupply at 4 d and 8 d. The WEGO plot describes the GO annotation and classification of DEGs, with the left *y*-axis showing the percentage of genes for the respective GO terms (red bars for down-regulated genes and grey bars for up-regulated genes) and the right *y*-axis depicting the number of both up- and down-regulated genes. Percentage and number of genes were calculated for the three broad main categories listed on the *x*-axis.

Enriched pathways represented by all DEGs at 4d and 8d of Fe deficiency

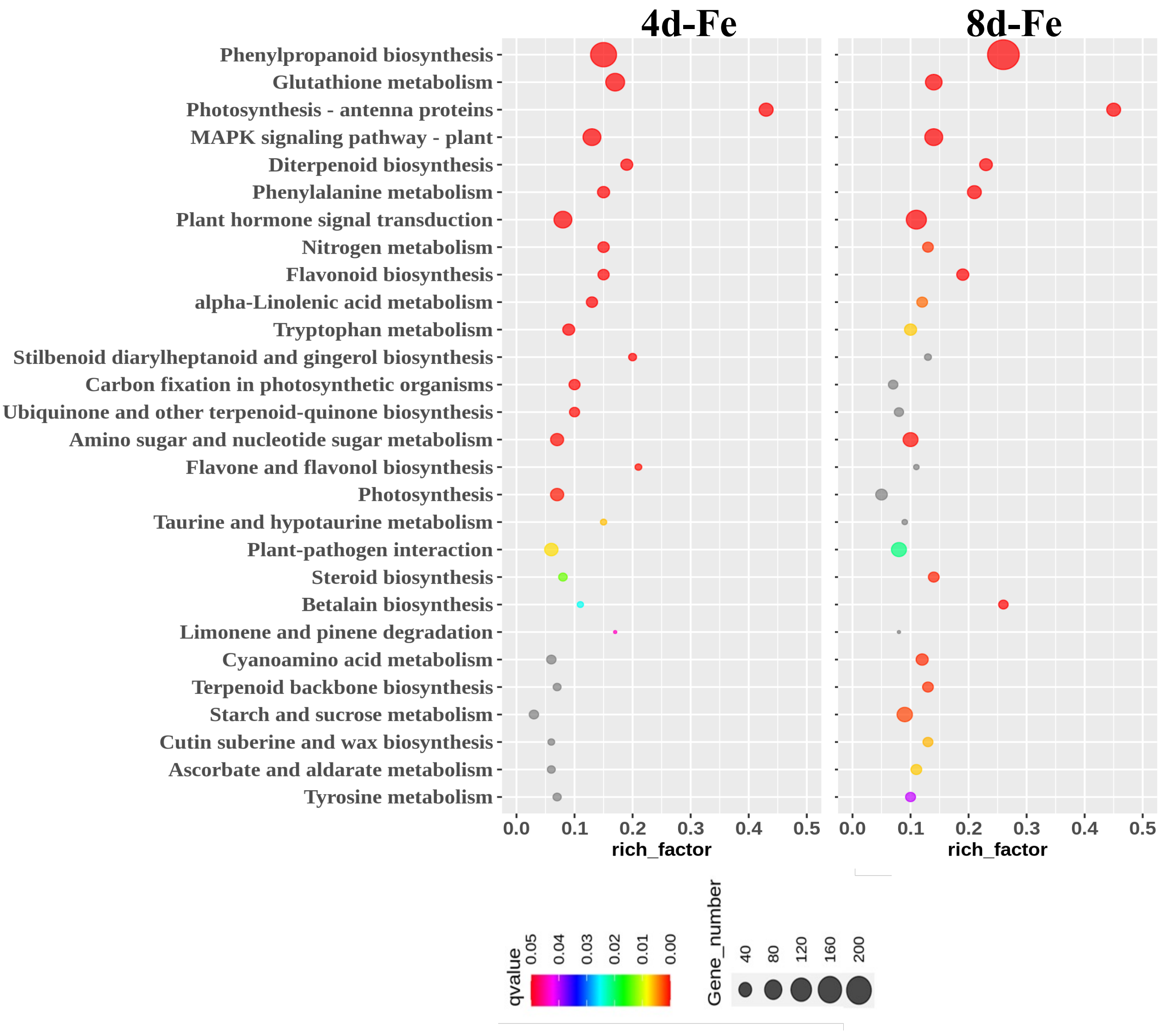


**Figure S5: KEGG pathway enrichment analysis (qvalue<0.05) for the genes showing altered expression in response to 4 d and 8 d of Fe deficiency w.r.t. control wheat roots.** X-axis represents the rich-factor (altered pathway genes/total pathway genes) and y-axis represents the pathway names. Bubble sizes are proportional to the number of perturbed genes in a pathway and colors represent significance levels as shown in the scale.

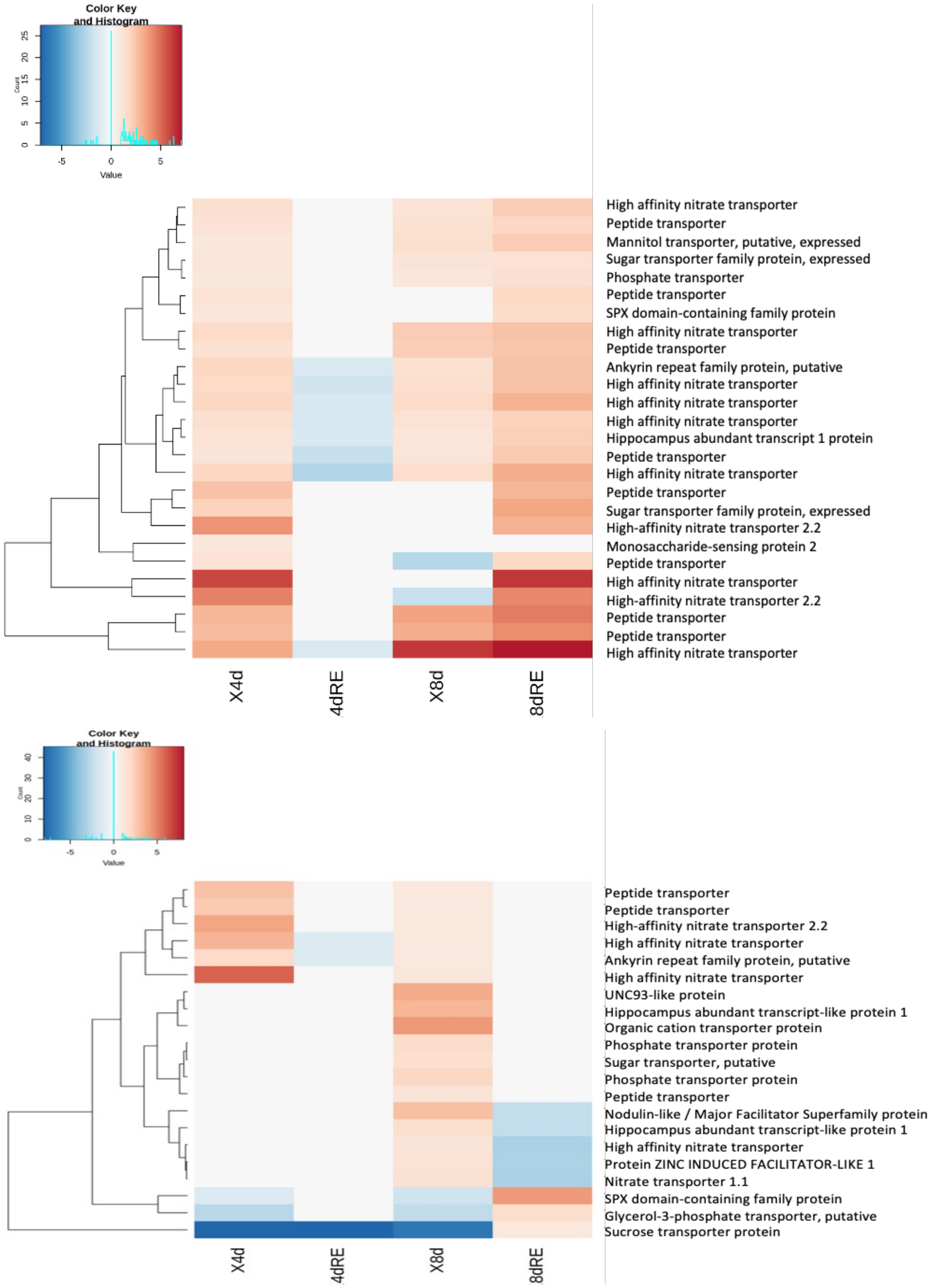


**Figure S6: Heatmaps depicting MFS genes that show an altered expression at 4 d (top panel) and 8 d (bottom panel) of Fe deficiency and further show a reverted expression response or no change in expression upon Fe resupply at the respective time-points.**

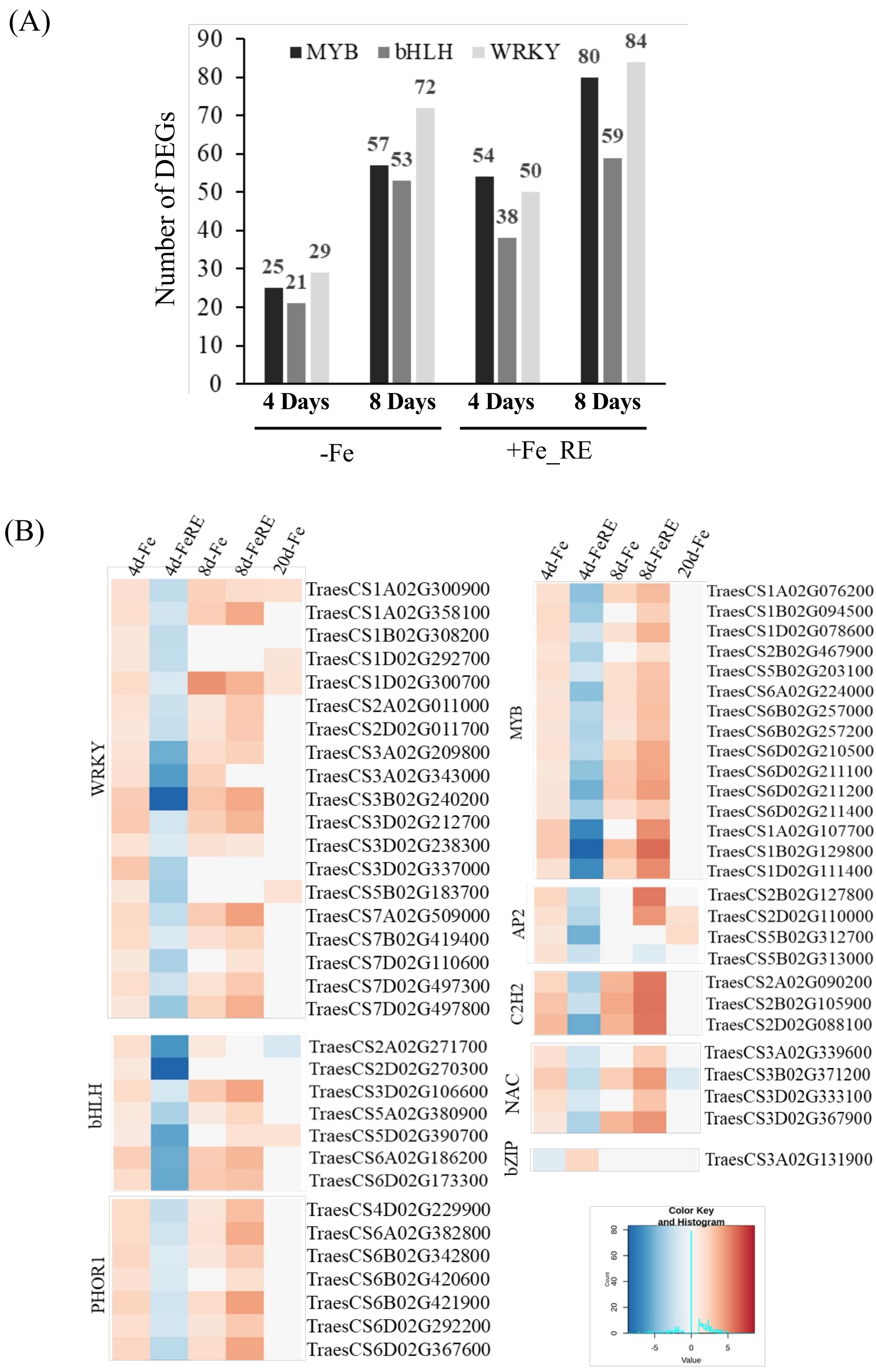


**Figure 7: Differential transcriptional response at 4 d, 8 d of Fe deficient conditions.** (A) Bar graph depicting number of altered Myb, bHLH and WRKY family TFs under Fe deficient as well as Fe resupply conditions at the respective time-points. (B) Heatmap analysis of major TF gene family representing those are altered under 4 d of Fe deficiency and show reverted expression patterns upon Fe resupply.

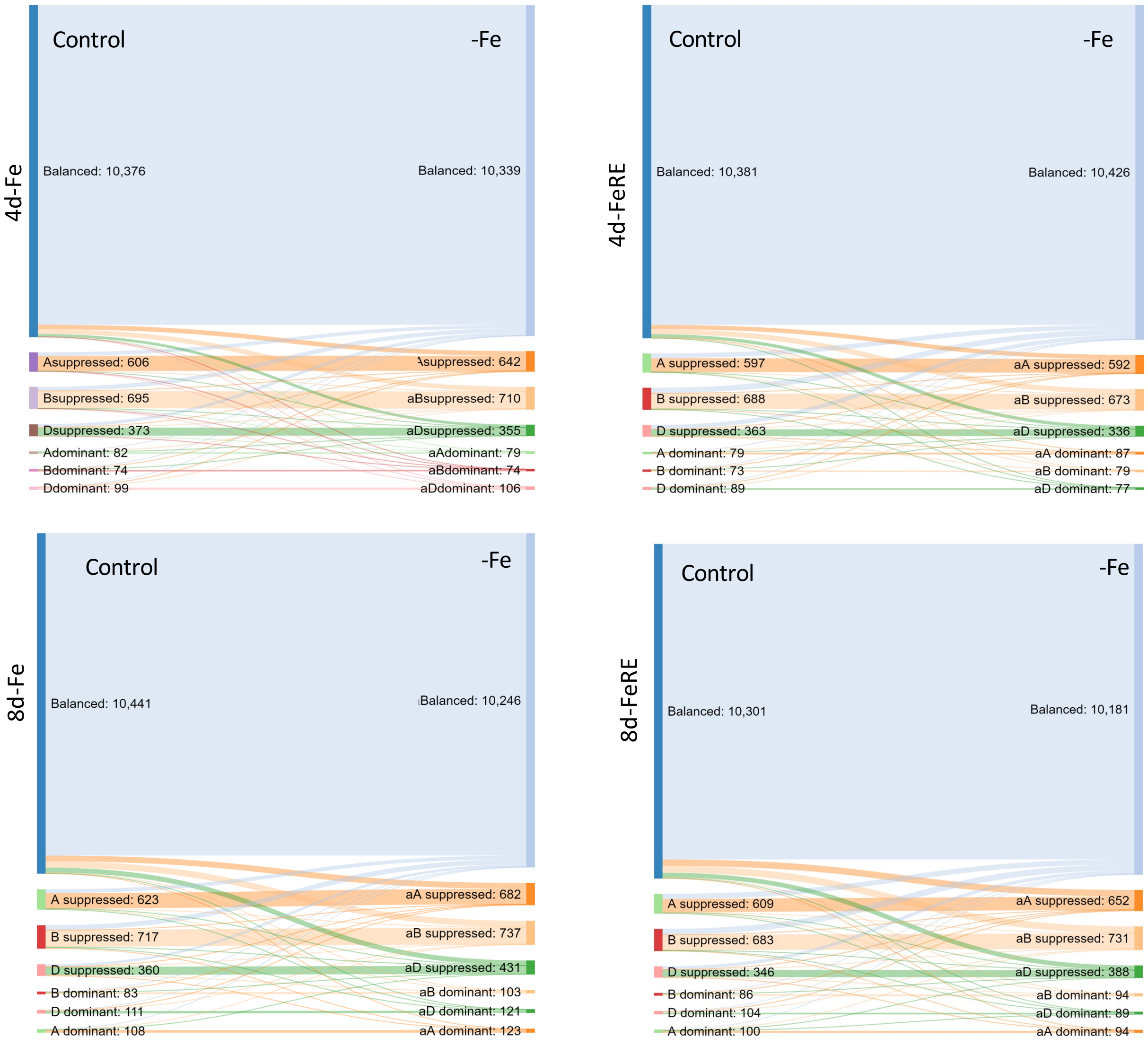


**Figure S8: Homoeolog expression bias in control and Fe deficient conditions depicted through sankey plots.** Distinct colours show the flow of triads from the seven defined categories from the control condition into the same category under –Fe or transition into another category as an effect of Fe deficiency.
